# Rare Ribosomal RNA Sequences from Archaea Stabilize the Bacterial Ribosome

**DOI:** 10.1101/2022.07.15.499945

**Authors:** Amos J. Nissley, Petar I. Penev, Zoe L. Watson, Jillian F. Banfield, Jamie H. D. Cate

**Affiliations:** Department of Chemistry, University of California, Berkeley, Berkeley, California, 94720, United States; Innovative Genomics Institute, University of California, Berkeley, Berkeley, California, 94704, United States; California Institute for Quantitative Biosciences, University of California, Berkeley, Berkeley, California, 94720, United States; Earth and Planetary Science, University of California, Berkeley, Berkeley, California, 94720, United States; Environmental Science, University of California, Berkeley, Berkeley, California, 94720, United States; Department of Molecular and Cell Biology, University of California, Berkeley, Berkeley, California, 94720, United States; Molecular Biophysics and Integrated Bioimaging Division, Lawrence Berkeley National Laboratory, Berkeley, California, 94720, United States

## Abstract

Ribosomes serve as the universally conserved translators of the genetic code into proteins and must support life across temperatures ranging from below freezing to above the boiling point of water. Ribosomes are capable of functioning across this wide range of temperatures even though the catalytic site for peptide bond formation, the peptidyl transferase center, is nearly universally conserved. Peptide bond formation by the ribosome requires correct positioning of the 3’
s-end of the aminoacylated tRNA (aa-tRNA) substrate, which is aided by an RNA hairpin in the ribosomal RNA (rRNA) of the large subunit, termed the A loop. Here we find that Thermoproteota, a phylum of thermophilic Archaea, substitute cytidine for uridine at large subunit rRNA positions 2554 and 2555 (*Escherichia coli* numbering) in the A loop, immediately adjacent to the binding site for the 3′-end of A-site tRNA. We show by cryo-EM that *E. coli* ribosomes with uridine to cytidine mutations at these positions retain the proper fold and post-transcriptional modification of the A loop. Additionally, these mutations do not exert a dominant negative effect on cellular growth, protect the large ribosomal subunit from thermal denaturation, and increase the mutational robustness of nucleotides in the peptidyl transferase center. This work identifies sequence variation in the peptidyl transferase center of the archaeal ribosome that likely confers stabilization of the ribosome at high temperatures and develops a stable mutant bacterial ribosome that can act as a scaffold for future ribosome engineering efforts.

## INTRODUCTION

Ribosomes are ribonucleoprotein complexes that carry out protein synthesis through the translation of genetic information into polypeptides. The ribosome consists of a large subunit which contains the catalytic peptidyl transferase center (PTC) and the small subunit where messenger RNA (mRNA) is decoded. Nucleotides in the PTC critical for peptide bond formation are universally or highly conserved, and mutation of these nucleotides often results in severe growth defects or lethal phenotypes.^1–3^ However, some mutational flexibility exists in the PTC beyond the core nucleotides that are directly involved in catalysis.^4^ Sequence variation at these sites, as well as post-transcriptional modification, are known to afford resistance to certain antibiotics targeting the ribosome.^5^ However, the functional impact of natural variation in the PTC on translation remains to be determined.

During translation elongation, an incoming aminoacyl transfer RNA (aa-tRNA) docks in the A site of the ribosome. For proper tRNA substrate positioning to occur, the tRNA anticodon must base pair with the mRNA in the small subunit decoding site while the tRNA 3’
s-CCA nucleotides interact with the A loop, an RNA hairpin in the ribosomal RNA (rRNA) of the large subunit.^6^ Proper orientation of the 3′-CCA end of the A-site tRNA by the A loop is necessary for correct positioning of the linked amino acid and subsequent peptide bond formation. In the *E. coli* ribosome, the interaction between the A-site tRNA 3′-CCA end and the A loop is formed in part by base pairing between universally conserved nucleotides C75 in the tRNA and G2553 in 23S rRNA (*E. coli* numbering).^7^ Mutation of either of these residues leads to a complete loss of activity that can be recovered by compensatory mutations that restore base pairing, establishing this interaction as vital to translation.^7,8^

While some sequence variability exists within the A loop, its structure is universally conserved due to its critical role in translation elongation.^9^ Additionally, 2’
s-*O*-methylation of U2552 is a conserved post-transcriptional modification that enables the proper folding of the A loop and ensures residues in the apical loop can interact with the A-site tRNA.^10^ In *E. coli*, loss of U2552 2′-*O*-methylation leads to severe growth defects and ribosome misfolding.^11^ Positions 2554 and 2555 in the A loop are conserved as uridines in eukaryotes and bacteria. U2554 forms a non-canonical base pair with C74 of the A-site tRNA and the base of U2555 stacks with C74.^12^ These interactions help to further stabilize the 3′-CCA end of the A-site tRNA upon docking in the ribosome. Previously, point mutations have been made at positions 2554 and 2555 in the bacterial ribosome with varying functional effects. U2555 mutations to adenine or guanine increase +1/-1 frameshifting and stop codon readthrough.^13^ In *in vitro* assembled ribosomes, a U2554C mutation decreased translation activity while a U2555C mutation had no effect.^4^ Despite some mildly deleterious effects, these studies demonstrate that there is some mutational flexibility at these positions.

Interestingly, positions 2554 and 2555 are sometimes found as cytidines in Archaea.^7,13^ It is known that thermophilic organisms stabilize their rRNA by increasing the GC content of rRNA, which stabilizes secondary structure.^14^ Additionally it has been shown that thermophilic organisms have higher levels of post-transcriptional rRNA modification, including *N*^4^-acetylcytidine^15^ and 2’
s-O-methylation^16^, and that the extent of these modifications is correlated with growth temperature. Here, we find that organisms in archaeal superphylum Crenarchaeota have cytidines at positions 2554 and 2555 in the A loop (*E. coli* numbering). Using archaeal sequences as a guide, we show that U to C mutations at positions 2554 and 2555 in 23S rRNA of the *E. coli* ribosome have a stabilizing effect on the 50S ribosomal subunit. This work identifies rare sequence variation in ribosomal RNA that likely leads to thermostability in the archaeal ribosome and describes how the specific stabilization of A loop structure can lead to global ribosome stabilization. These insights into how thermophilic organisms stabilize their ribosomes provide a foundation for engineering ribosomes with increased mutational robustness and evolvability.

## RESULTS

### Sequence Conservation in the A loop

To investigate sequence diversity in the A loop, we aligned archaeal 23S rRNA sequences from the SILVA database^17^ to a structurally guided sequence alignment of large subunit (LSU) rRNA^9^, identifying nucleotide variation at positions 2554 and 2555 (*E. coli* numbering) (**Figure 1A**). Both positions are almost universally uridine residues (UU), except for sequences from the Crenarchaeota superphyla (**Figure 1B**). Within this superphyla, the phylum Nitrososphaerota generally includes mesophilic organisms that grow at ambient temperatures, while Thermoproteota includes hyperthermophilic organisms that grow at high temperatures.^18^ We find that Nitrososphaerota tend to have a cytidine at position 2554 (CU) while Thermoproteota more often have cytidines at both positions (CC) (**Figure 1**). Organisms within Thermoproteota have extreme growth temperatures even among thermophiles, suggesting that this change may play a role in ribosome stability. However, there are some hyperthermophiles in Crenarchaeota that have either CU or UU sequences in the A loop. Additionally, other thermophilic Archaea, such as organisms in Thermoplasmatota and Euryarchaeota, have uridines instead of cytidines at these positions, suggesting that the CC mutation is one of multiple evolutionary strategies to confer heat resistance.

**Figure 1.**
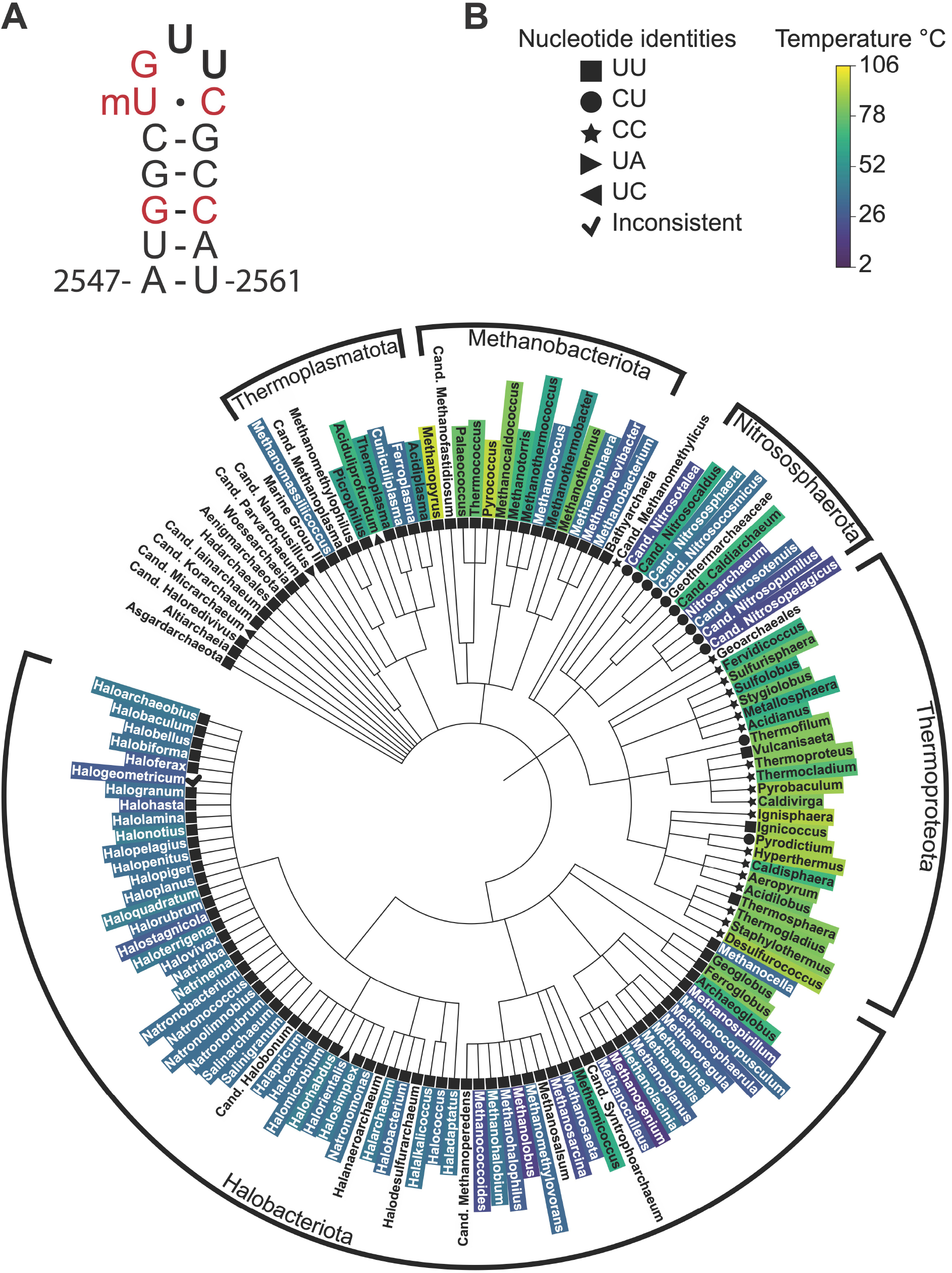
Distribution of nucleotide sequence variation in the A loop across Archaea. A) RNA secondary structure of the *E. coli* A loop. Red nucleotides are universally conserved. Positions 2554 and 2555 are in bold. B) Phylogenetic tree of archaeal large ribosomal subunit (LSU) rRNA in the SILVA database.^17^ Coloring at the genus level is based on optimal growth temperatures. Nucleotide sequences at the equivalent positions to 2554 and 2555 in *E. coli* 23S rRNA are indicated by geometric symbols. CU and CC variations at these positions are found mainly in Crenarchaeota (Nitrososphaerota and Thermoproteota).

### A loop Mutations in the *E. coli* 50S Ribosomal Subunit

Since nucleotides 2554 and 2555 in 23S rRNA are conserved as uridines in bacteria and eukaryotes, we studied the effects of U to C mutations, inspired by the Archaeal A loop sequences, in the context of the *E. coli* ribosome. We first assessed whether mutations in the A loop have a dominant negative effect on bacterial growth. *E. coli* were transformed with an IPTG-inducible pLK35 plasmid harboring either a wild-type (WT) 23S rRNA sequence, a U2554C (CU) point mutation, or a U2554C-U2555C (CC) double mutation. The growth of these transformants was then measured in the presence of IPTG (**Figure 2A**). All transformants grew with similar doubling times (**Table S1**). Because genomically encoded WT ribosomes are also present in the cell during the experiment, we cannot rule out that the A loop mutations affect *in vivo* translation. However, the plasmid-encoded tagged ribosomes comprise around half of cellular ribosomes at the end of the growth experiment (**Figure S1**). PTC mutations can have a dominant effect on cellular growth even in the presence of genomically encoded WT ribosomes.^3^ Since we do not see decreases in the growth rates for transformants, this indicates that the A loop mutations do not cause dominant growth defects.

**Figure 2.**
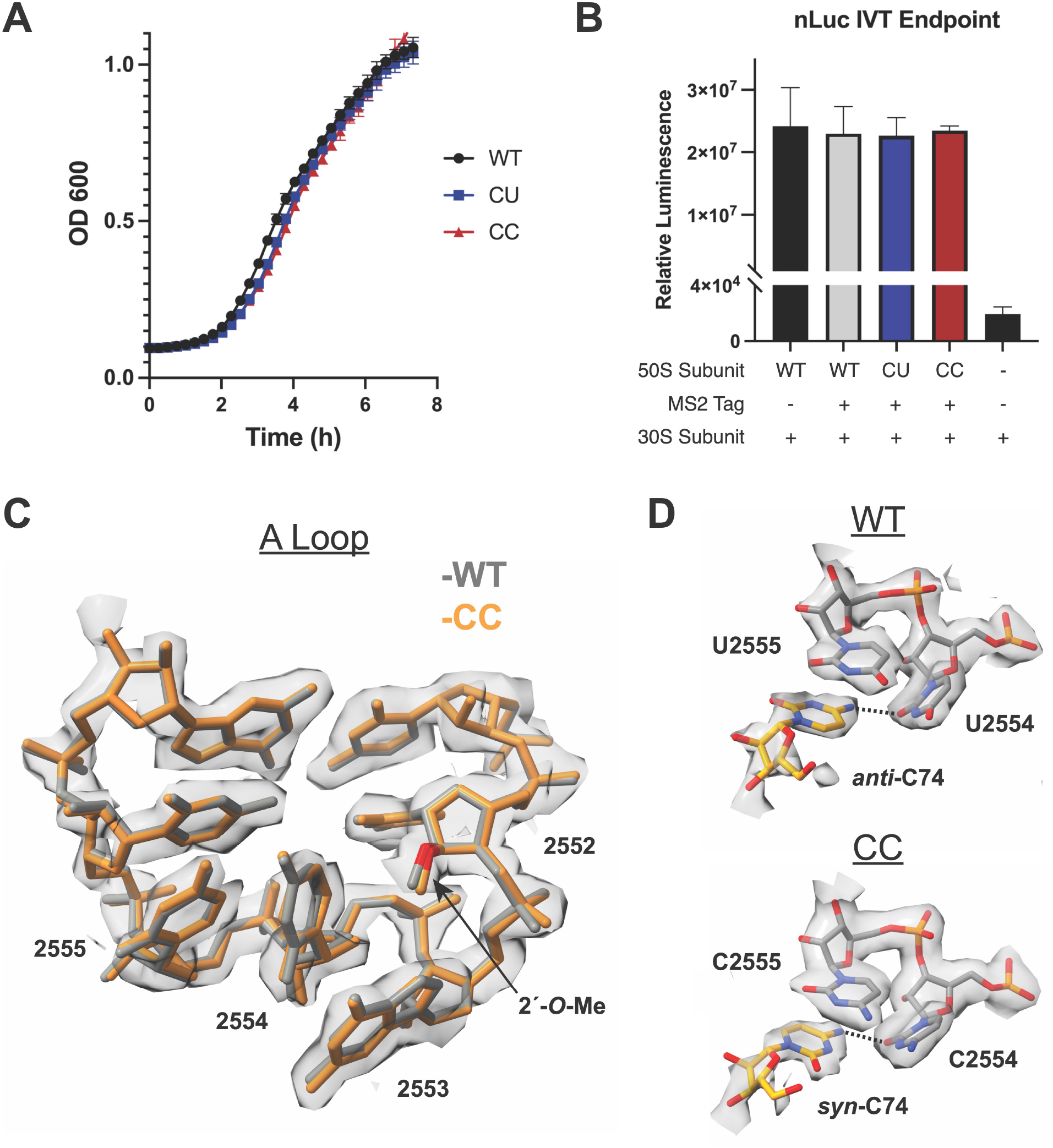
Effects of CU and CC mutations on the A loop. A) Growth curves of *E. coli* expressing plasmid-encoded ribosomes. The plasmid-encoded 23S rRNA harbors an MS2 tag. B) nLuc *in vitro* translation endpoint assay for mutant ribosomes at 37 °C. C) Cryo-EM density and model for the CC mutant ribosome are shown in grey and orange respectively. The model from a WT ribosome structure^21^ is shown in grey. D) (Top) Cryo-EM density and model for C74 in the A-site tRNA and U2554 and U2555 in the A loop from^21^. (Bottom) Cryo-EM density and model for C74 in the A-site tRNA and C2554 and C2555 in the A loop in the CC mutant ribosome. The C74 (N4)-U2554 or C2554 (O2) hydrogen bonds are represented by dashed lines. A *B*-factor of 30 Å^2^ was applied to the CC mutant ribosome cryo-EM map.

Having established the tolerance of U to C A-loop mutations *in vivo*, we further characterized the mutational effects *in vitro*. Three forms of the *E. coli* 50S ribosomal subunit were purified using an MS2 tag^19^: WT, U2554C (CU) and U2554C-U2555C (CC) (**Figure S2**). We first examined whether these mutations influence bacterial ribosome activity at normal growth temperatures using an *in vitro* translation endpoint assay in which full length nLuc protein (171 amino acids) is translated. At 37 °C, both the CU and CC ribosomes produce similar levels of active nLuc protein compared to WT ribosomes (**Figure 2B**), indicating that the U to C mutations do not cause translation defects *in vitro*.

### Structure of the CC 70S Ribosome

To probe the structural effects of the A loop mutants, we solved the cryo-EM structure of the CC mutant 70S ribosome in a complex with mRNA and aminoacyl-tRNAs in the P site and A site (non-hydrolyzable Met-tRNA^fMet^, see methods) to a global resolution of 2.2 Å (**Figure S3**). The structure reveals that, in the presence of the U to C mutations at positions 2554 and 2555, the A loop fold is indistinguishable from the WT A loop (**Figure 2C**). This result is in line with the observations that the CC ribosome has WT (UU) levels of activity at 37 °C *in vitro* and has no impact on cellular growth. In addition, the A loop in the CC mutant also contains the proper 2’
s-O-methylation of U2552, as evidenced by clear density for the 2′-*O*-methyl group on the ribose of U2552 (**Figure 2C**). Mutations at positions 2554 and 2555 thus do not affect the 2′-*O*-methylation of the nearby residue U2552 by *E. coli* methyltransferase RrmJ. The proper fold of the A loop in the CC structure is also indicative of the correct modification, as loss of U2552 2′-*O*-methylation has been shown to change the fold of the A loop.^10^

While the structure of the A loop itself is not altered by the CC mutations, the positioning of the A-site tRNA and its interaction with the A loop is altered in two ways compared to WT ribosomes. First, C74 of the A-site tRNA adopts a *syn* conformation, in contrast to the *anti*-conformation observed in the WT ribosome (**Figure 2D**).^20,21^ The *syn* conformation is highly disfavored thermodynamically for pyrimidines and therefore, *syn* pyrimidines are not frequently found in RNA.^22^ In the WT ribosome, C74 of the A-site tRNA is held in its position in part through hydrogen bonding between the exocyclic amine of C74 (N4) and the O2 carbonyl oxygen of U2554 (O2) in the A loop and by base stacking with U2555 in the A loop and C75 in the tRNA. In the CC mutant ribosome, hydrogen bonding between N4 of C74 and the O2 of C2554 is preserved with a *syn*-C74 (**Figure 2D**). No new hydrogen bonds are formed that stabilize this conformation and no conformational change in the ribosome occurs that would disfavor the *anti*-conformation. It is possible that either a change in the local electronic environment in the CC mutant or transient solvent interactions work to stabilize the *syn* conformation of C74. Despite this change, the C75-G2553 base pair is preserved and A76 and the linked methionine are in their canonical positions.^20,21^

Second, during cryo-EM data processing, a masked classification of the A site yielded two slightly shifted tRNA classes that were equally populated (**Figure S4**), one of which had clear density for the 3’
s-CCA end of the A-site tRNA that was further processed to yield the final high-resolution structure. The second A-site tRNA class identified in the cryo-EM data (class 2) lacks density for most of the tRNA acceptor stem (including the 3′-CCA) but contains density for the anticodon and weak density for the rest of the tRNA (**Figure S5**). This contrasts with the WT *E. coli* ribosome structure, where all A-site tRNA classes had density for the CCA-end.^21^ The structure of the WT ribosome used for comparison contains the same tRNAs in the P-site and A-site as in the CC ribosome structure, and both structures used paromomycin to favor A-site tRNA binding, which facilitates the direct comparison (methods). These results suggest that the U to C mutations in the A loop interfere with docking of the A-site tRNA’s 3′-CCA end, possibly due to the *syn*-C74 conformation favored by the mutation.

### CC Mutations Stabilize the 50S Ribosomal Subunit

Since CC variation in the A loop is found in hyperthermophilic Archaea, we sought to assess whether it stabilizes the 50S ribosomal subunit. To further examine ribosome activity in this context, we used a Nanoluc (nLuc) luciferase complementation assay, in which a truncated and catalytically inactive form of nLuc, termed 11S, is complemented by an *in vitro* translated peptide with high affinity for 11S, termed HiBit, to restore luciferase activity (**Figure 3A**).^23^ This assay enables the close monitoring of translation rates, rather than the endpoint detection of translated protein as in the nLuc assay, and is able to detect low levels of ribosome activity. We find that ribosomes with U to C mutations at positions 2554 and 2555 are as active as WT ribosomes at 37 °C in the HiBit assay (**Figure 3A**), which is consistent with the nLuc endpoint assay (**Figure 2B**).

**Figure 3.**
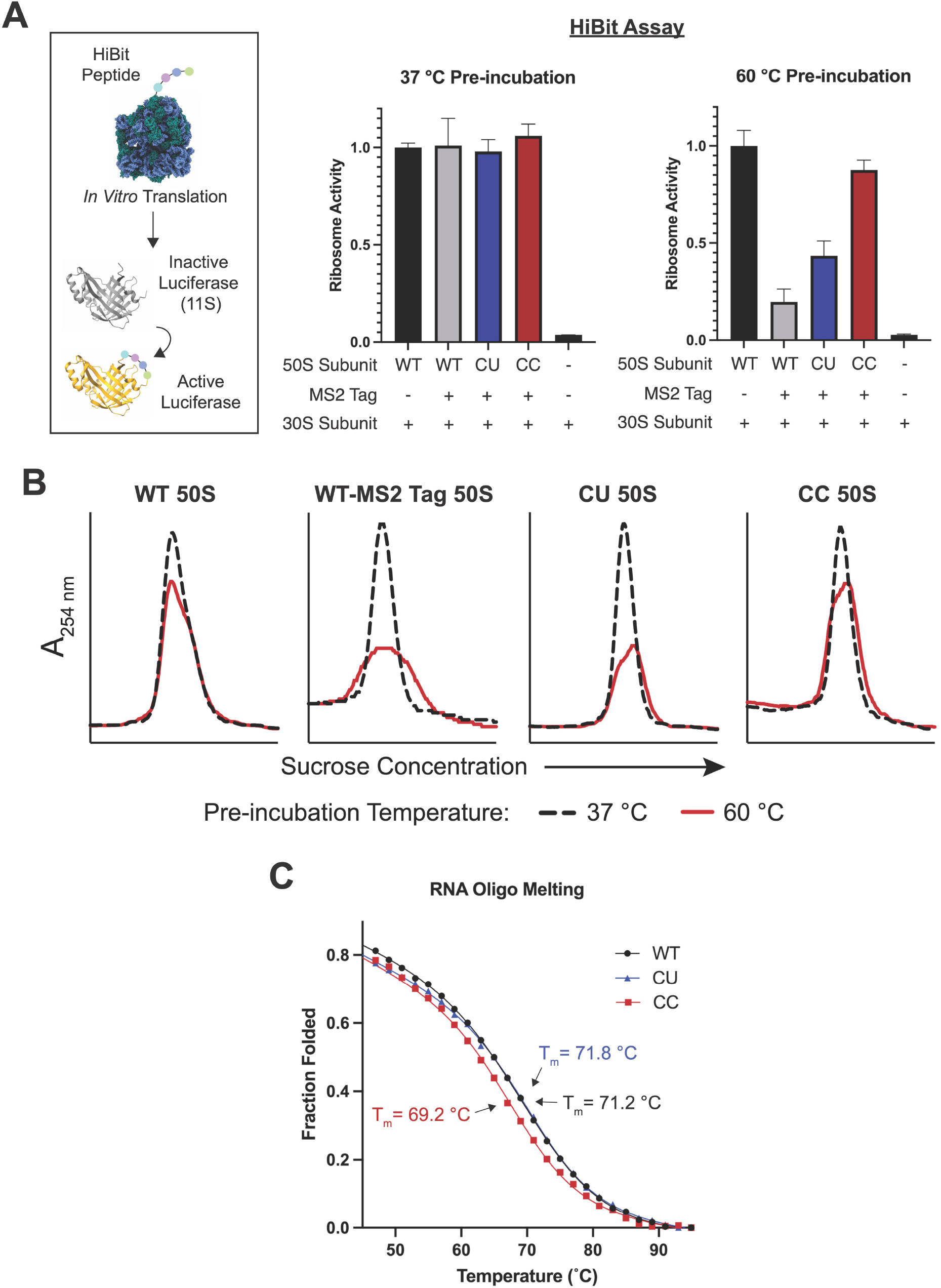
Stability of 50S subunits with A loop mutations at positions 2554-2555. A) In the HiBit assay, the 12 amino acid HiBit peptide is translated by the ribosome, which activates a truncated luciferase (11S). 50S subunits were pre-incubated at 37 °C or 60 °C and then their relative activities were determined with the HiBit assay. Both untagged and MS2-tagged WT 50S subunits were used as controls. B) 50S subunits were heat treated at 37 °C (black dash) or 60 °C (red) and then resolved on a 15-40% sucrose gradient. C) RNA oligonucleotides (15 nucleotides) containing the WT A loop sequence (2547-2561), the CU, or CC mutant sequences were melted on a CD spectrometer. The melting temperature (T_m_) was calculated for each construct using a two-state model.

*E. coli* protein components of an *in vitro* translation reaction are not thermostable^24^, and rather than conducting *in vitro* translation assays at temperatures above 37 °C, we sought to decouple the thermostability of the 50S subunit and the translation reaction mixture. To do this, 50S subunits were first incubated at high temperatures, cooled slowly, and then added to *in vitro* translation reactions. Using this experimental design, we tested the thermostability of 50S subunits with the HiBit assay. WT ribosome activity begins to decrease after incubation at 60 °C, which suggests that this is the temperature at which 50S subunit denaturation begins (**Figure S6**). Surprisingly, we found that the MS2 tag inserted into 23S rRNA helix 98 (H98) affects ribosome stability, as the tagged WT ribosome has considerably lower levels of activity than the untagged WT ribosome after 50S subunit incubation at 60 °C (**Figure 3A** and **Figure S6**). MS2 tags in H98 have been used extensively as they do not affect cellular growth or ribosome assembly and function.^19,25,26^ In further experiments, we sought to asses if CC mutations could recover activity of MS2-tagged 50S subunits after incubation at 60 °C.

Comparison of the MS2-tagged WT ribosomes and the tagged mutant ribosomes shows that both the CU and CC mutants have higher levels of activity than the tagged WT ribosome after 60 °C incubation of the 50S subunit, with the CC mutant retaining the highest levels of activity (**Figure 3A**). The higher activity retained by the CC ribosome compared to WT suggest that these mutations stabilize the ribosome. Notably, as in the HiBit assay, we also see increased protein output from the CC mutant compared to the tagged WT 50S subunits after pre-incubation at 60 °C in a full length nLuc endpoint assay (**Figure S7**).

To assess whether the effects on ribosome activity observed in the HiBit assay were due to possible folding defects in the PTC after heat treatment, or due to global misfolding of the 50S subunit, 50S subunits were resolved on sucrose gradients. Consistent with the HiBit assay, untagged WT 50S subunits are well folded after heat treatment at 60 °C with only a slight loss of folded 50S subunits (**Figure 3B**). The addition of an MS2 tag into H98 (WT-MS2 Tag 50S) leads to global ribosome unfolding after heat treatment. The peak height of the tagged WT 50S is decreased and broader compared to the untagged 50S, consistent with a heterogeneous population of 50S subunits, likely due to degradation. While the CU 50S subunit shows slightly higher activity in the HiBit assay than the tagged WT subunit, the CU mutation does not improve global 50S subunit stability substantially, as determined with sucrose gradients (**Figure 3B**). This matches the full length nLuc endpoint assay where the CU subunit has low activity after heat treatment (**Figure S7**). In contrast, as demonstrated with *in vitro* translation assays, the CC 50S subunits remain more homogeneous on sucrose gradients, indicating they are better folded after heat treatment than the tagged WT subunit and that CC mutations globally stabilize the 50S subunit.

To further probe the basis for ribosomal stabilization by U to C mutations in the A loop, we conducted thermal melting experiments of isolated rRNA hairpins comprising the A loop. We used RNA oligonucleotides of WT and mutant A loop sequences (CU and CC), all of which contained 2’
s-O-methylation of U2552. In circular dichroism (CD) thermal melting experiments, the RNA constructs demonstrated cooperative and reversible unfolding (**Figure S8**). The WT, CU, and CC A loop constructs had melting temperature of 71.2 °C, 71.8 °C, and 69 °C, respectively (**Figure 3C** and **Table S2**) in low ionic conditions (methods). These high melting temperatures suggest that the A loop may retain its secondary structure in the ribosome at the high temperatures used for the heat treatment experiment, regardless of whether nucleotides 2554-2555 are UU, CU, or CC. These results also suggest that the CU and CC mutations may only stabilize the structure of the terminal loop of the A loop hairpin in the context of the intact ribosome.

### Additional Mutations at the Base of the A loop RNA Hairpin

In view of the global ribosomal stabilization conferred by CC mutations in the A loop, we also searched archaeal rRNA sequences for additional A loop sequence variation at other positions in the RNA hairpin. We found that most Archaea with the CC variation have a GC base pair between nucleotides 2548 and 2560 (base pair 2) (**Figure 4A** and **Figure S9**). This GC base pair is also present in over half of Archaea with uridines at positions 2554 and 2555, whereas *E. coli* has a less stable AU base pair at this position. As thermophilic organisms have increased rRNA GC content^14^, we hypothesized that secondary structure variation at the base of the A loop might also serve to stabilize the ribosome. The base of the A-loop RNA helix interacts with ribosomal protein uL14, which varies significantly between Bacteria and Archaea in this region and complicates incorporation of an Archaeal A loop helical sequence into the *E. coli* ribosome. Because of this, we also looked at the A loop of the thermophilic bacterium *Thermus thermophilus*. The *T. thermophilus* A loop also has a GC base pair at base pair 2, and the uL14-rRNA contacts in *E. coli* and *T. thermophilus* are more similar.^20^

**Figure 4.**
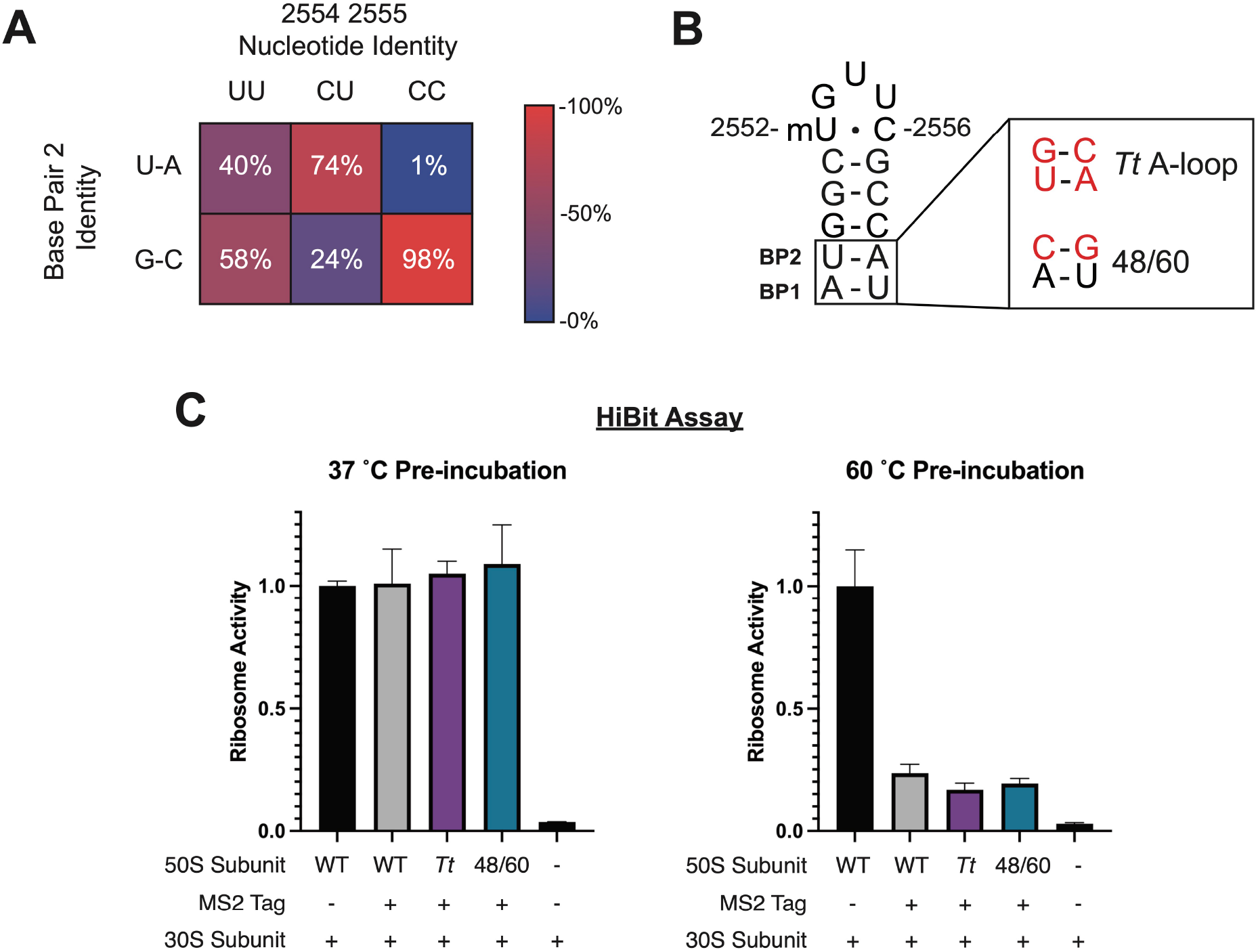
Effects of A-loop secondary structure variation on global 50S subunit stability. A) Distribution of archaeal 23S sequences from the SILVA database^17^ based on nucleotide identity at positions 2554 and 2555 (top) and the identity of the base pair at position 2 (2548-2560) (left). The heatmap is colored to represent the percentage of each column that has either a U-A or G-C base pair. B) The helix of the *E. coli* A-loop was mutated to match that of *T. thermophilus* (top). A second A loop mutant (48/60) was designed to maintain interactions between the A-loop RNA stem and *E. coli* ribosomal protein uL14. C) 50S subunits were pre-incubated at 37 °C or 60 °C and then their relative activities were determined with the HiBit assay.

Based on Archaeal and *T. thermophilus* A-loop sequences, we designed two mutant *E. coli* 50S ribosomal subunits to test for global stability. The first incorporates the A-loop sequence from *T. thermophilus* (*Tt* A loop), which swaps the order of the A and U at base pair 1 and changes base pair 2 to a GC base pair. These changes are predicted to stabilize the secondary structure of the A loop hairpin (**Figure S10**). To maintain native uL14 interactions with the A loop, a second ribosome mutant (48/60) was prepared. With this design, base pair 2 of the A loop is changed to a GC base pair but pyrimidine bases are kept at positions 2548 and 2561 (**Figure 4B**). *E. coli* uL14 interacts with the pyrimidine O2 at those positions and the 48/60 mutant should preserve the native contacts. While the 48/60 mutant is predicted to be less stable than the Tt A loop mutant, it is predicted to afford more stability than the WT *E. coli* sequence (**Figure S10**). We assayed the thermostability of these mutants using the HiBit assay. As expected, these mutant ribosomes have full activity after pre-incubation at 37 °C indicating that the mutations do not affect the fold of the A loop (**Figure 4C**). After pre-incubation of the 50S subunits at 60 °C, the *Tt* A loop and 48/60 mutant ribosomes exhibit low activity, similar to that of the MS2-tagged WT ribosome control. These results indicate that these mutations in the RNA helix adjacent to the A loop do not confer additional stability to the WT ribosome on their own, in contrast to the stability afforded by U to C mutations in the apical loop.

### CC Mutations Increase the Mutational Robustness of PTC Nucleotides

Stabilization of the PTC could have utility in ribosome engineering where mutations are made in and around the PTC to allow for the incorporation of nonnatural amino acids or other monomers into proteins.^27,28^ As CC mutations stabilize the 50S subunit from thermal denaturation, we posited that they could stabilize the PTC in the presence of other mutations. To investigate whether CC mutations increase the mutational robustness of the PTC, 50S ribosomal subunits carrying an A2451C point mutation or a U2554C-U2555C-A2451C mutation (CC-A2451C) were purified using the MS2-tag purification. The A2451C point mutation was chosen because A2451 is a core PTC nucleotide involved in catalysis and mutations at this position have been shown to greatly decrease the rate of peptide bond formation.^29^ Additionally, A2451 is over 20 Å from the A loop (**Figure 5A**) and any observed stabilization from the CC mutations would have to come from global stabilization of the PTC rather than local short range interactions.

**Figure 5.**
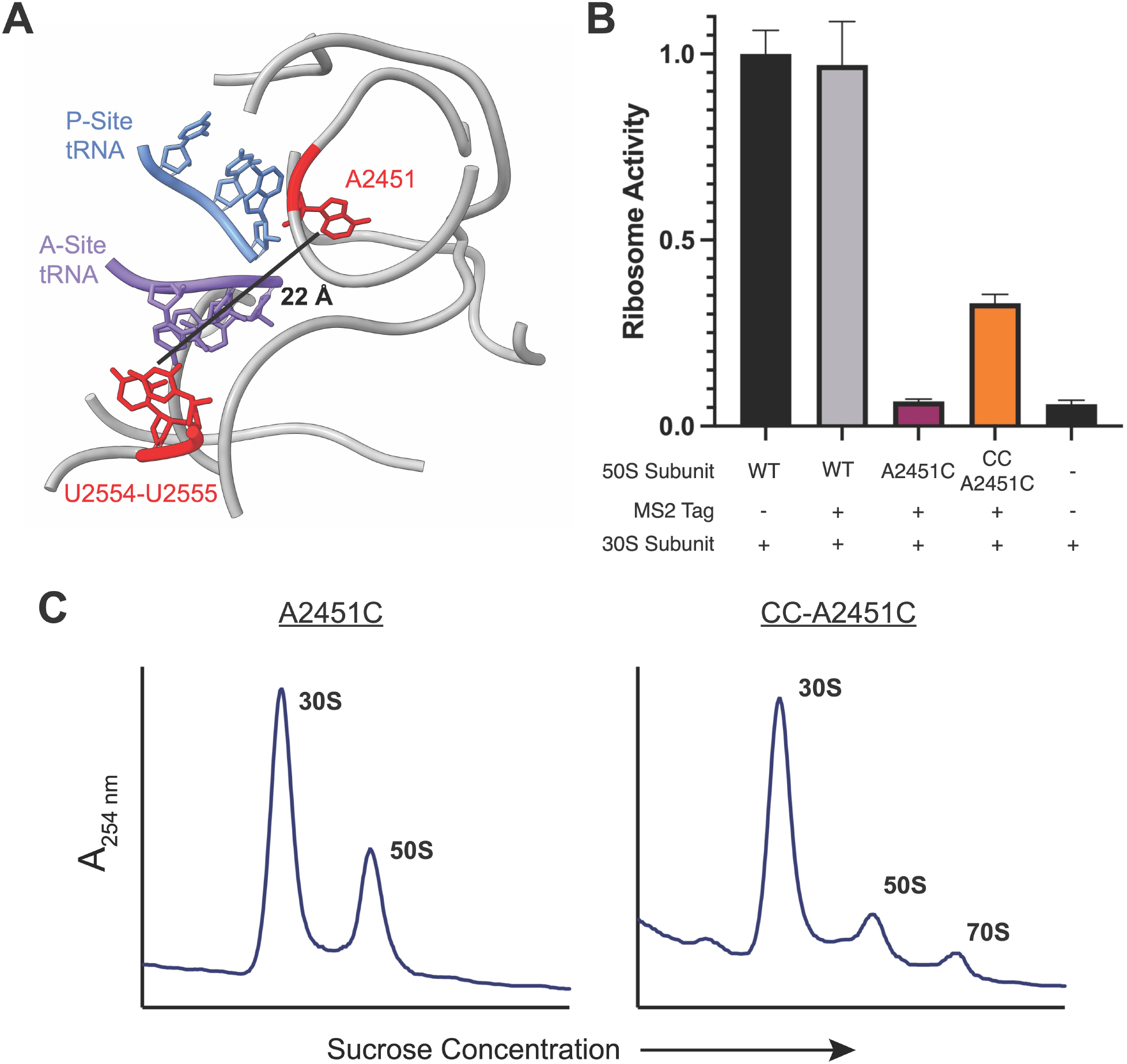
Mutational robustness of the PTC in CC ribosomes. A) Model of the *E. coli* PTC (PDB:7K00)^39^. The CCA-ends of the P-site (blue) and A-site (purple) tRNAs are highlighted along with 23S rRNA bases U2554, U2555, and A2451C (red). The distance between U2555 and A2451C is marked. B) The relative activities of 50S subunits with an A2451C mutation or the triple mutation CC-A2451C were determined using the HiBit assay. C) 70S formation by A2451C (left) and CC-A2451C 50S subunits (right).

We used the HiBit assay to determine the activities of ribosomes with A2451C and CC-A2451C mutations at 37 °C (**Figure 5B**). A2451C ribosomes have very low activity–close to the background signal in the HiBit assay–indicative of a severe defect in translation. The addition of CC mutations to A2451C yields a more active ribosome, suggesting that CC mutations can compensate for some of the PTC misfolding caused by an A2451C mutation. It is worth noting that the observed increase in activity cannot be due to WT ribosome contamination as the purified CC-A2451C 50S sample has less contamination (∼2% contamination) than the A2451C 50S sample (∼5% contamination) (**Figure S11**). In line with its low activity, sucrose gradient traces indicate that A2451C 50S subunits do not form 70S ribosomes *in vitro*, while the CC-A2451C 50S subunits retain some ability to form 70S ribosomes (**Figure 5C**). Together, these data indicate that CC mutations can compensate for the destabilizing effects of deleterious PTC mutations.

## DISCUSSION

While the ribosome is highly conserved and contains universally conserved elements across all domains of life, hyperthermophiles have adapted their ribosomes to function at extreme temperatures. Here we find that the hyperthermophilic Archaea of the Thermoproteota phylum have adapted rare rRNA sequence variation in the catalytic center of the ribosome, in which cytidines are found at positions 2554 and 2555 in the A loop of 23S rRNA. Grafting U to C mutations at positions 2554 and 2555 in *E. coli* 23S rRNA stabilizes the 50S ribosomal subunit from thermal inactivation, consistent with CC variation serving a stabilizing role in certain hyperthermophilic Archaea. Consistent with our findings, some organisms in Thermoproteota have recently been shown to stabilize their tRNA through 2’
s phosphorylation of a uridine residue, which highlights the need for stabilized translation machinery at high temperature.^30^

CC sequence variation is only observed in the Thermoproteota phylum. This can be explained phylogenetically if the common ancestor of Nitrososphaerota and Thermoproteota developed the CC modification. This modification was advantageous for Thermoproteota as thermophilic organisms and was therefore evolutionary stable at high growth temperatures. However, this modification may have slowly reverted in the mesophilic Nitrososphaerota to CU. It is notable that not all thermophilic Archaea and no thermophiles in the other domains of life acquired this variation. This suggests that there may be an evolutionary disadvantage of the CU and CC variation.

The cryo-EM structure of the CC mutant ribosome at 2.2 Å resolution revealed an unexpected change in the interaction between the mutant A loop and the A-site tRNA. In contrast to UU ribosomes, nucleotide C74 in the tRNA adopts the disfavored *syn* conformation when bound to the CC mutant ribosome, which may be a side effect of a change in the local electronic environment by the U to C mutations. *Syn* pyrimidines are exceptionally rare and are more commonly found in the aptamer binding domains and catalytic sites of RNAs.^22^ The *syn*-C74 in the A-site tRNA, at lower temperatures such as those tested with the *E. coli* ribosome, may lead to slight translation defects, which we were unable to detect in bulk translation assays, and could explain why CC A loop mutations are not more prevalent. Additionally, it is possible that the *syn* conformation is less populated at the high growth temperatures of Thermoproteota and therefore less problematic for hyperthermophiles. During cryo-EM processing we identified a highly populated A-site tRNA class that has a disordered acceptor stem. This may indicate that the CC ribosome is deficient in A-site tRNA binding. In yeast, a pseudouridine to cytidine mutation at nucleotide 2554 (*E. coli* numbering) decreased tRNA binding to the ribosome.^31^ It is also possible that a *syn*-C74 may interfere with tRNA translocation after peptide bond formation. When tRNA docks in the ribosomal P site, C74 directly base pairs with G2252 in the large subunit rRNA P loop. This base pair would require the *syn*-C74 in the A site tRNA to flip back to the *anti*-conformation when it translocates to the P site, which could present a hurdle for proper P-site docking. However, tRNAs with mutations at position 74 are still accepted as substrates by the ribosome, which indicates that the C74-G2552 interaction is not essential for translation.^8,32^ This suggests that a tRNA containing a *syn*-C74 could be translocated by the ribosome with little difficulty.

It is striking that modest pyrimidine to pyrimidine substitutions at only two positions can globally stabilize the 50S ribosomal subunit, a 1.5 MDa particle. As CD melting experiments demonstrate that the secondary structure of the RNA hairpin containing the A loop is not stabilized by U to C mutations, the observed stabilization of the mutant 50S ribosomal subunit likely arises from enhanced tertiary interactions in the A loop that only form within the ribosome. This is consistent with a prior structural analysis of an isolated A loop RNA hairpin, which differs from the structure of the A loop within the 50S ribosomal subunit, especially in the apical loop.^33^ The A loop rRNA helix has a high melting temperature outside of the ribosome (71.2 °C), which suggests that the A loop helix may stay folded in the ribosome even at high temperatures. Additionally, mutations in the A loop predicted to stabilize the secondary structure of the rRNA helix had no additional impact on the global thermal stability of the 50S subunit. Taken together, these data suggest that the stability of A-loop secondary structure may not be as important for global ribosome stability as its tertiary structure. As the A loop is part of the last region of 23S rRNA to fold^34^, its tertiary stabilization may lead to global stabilization of the 50S ribosomal subunit.

Protein engineering can benefit from a thermostable scaffold which can better withstand the incorporation of destabilizing mutations that are needed to evolve new functionality.^35^ For example, chimeric proteins which incorporate protein domains from thermophilic archaea have been used to improve engineering of *E. coli* aminoacyl-tRNA synthetases.^36^ Here, we show that rare sequence variation in highly-conserved regions of rRNA from thermophilic organisms can stabilize the *E. coli* ribosome. The CC mutant *E. coli* ribosome does not have an observable defect *in vivo* and is able to translate both small peptides and full proteins *in vitro*. Previous computationally designed ribosomes with mutations in the A loop had severe defects in ribosome activity.^37^ In contrast, the mutations examined here, including those in the stem of H92, preserve conserved nucleotides and lead to highly functional ribosomes. While mutating the PTC has allowed for the incorporation of non-proteinogenic monomers by the ribosome, PTC mutations can have a severely destabilizing effect.^4,38^ We show that CC mutations stabilize the ribosome at high temperatures in the context of a destabilizing MS2 tag in the 50S subunit and that they can increase the activity of a ribosome with a mutation of a critical PTC nucleotide. The CC mutations in the A loop may therefore increase the mutational robustness and evolvability of the surrounding PTC and enable future engineering of ribosomes capable of improved synthesis of non-proteinogenic polymers.

## MATERIAL AND METHODS

### Archaeal rRNA Sequence Alignment and Tree Visualization

To investigate sequences within the A loop we aligned archaeal 23S rRNA sequences from the SILVA database^17^ to a manually curated sequence alignment of large subunit (LSU) rRNA^9^ using mafft with the option –add.^40,41^ We used a custom python script (https://github.com/petaripenev/A-loop_sequence_analysis) to identify relevant positions, and correlate data for average optimal living temperature from the TEMPURA database^42^ with the alignment and a phylogenetic tree retrieved from SILVA. For several new Nitrososphaerota taxa missing from the TEMPURA database, we searched published articles to identify likely ranges of optimal living temperature (**Table S3**) The nucleotide identity and average optimal living temperature were mapped on the tree with iTOL.^43^ Phyla were indicated following the latest GTDB taxonomy.^44^

### Cloning and Plasmid Design

A modified version of the pLK35^45^ plasmid, which encodes 5S, 16S, and MS2-tagged 23S rRNA^19^, with an IPTG inducible tac promoter was used for ribosome mutagenesis. Point mutations were introduced using the Q5 mutagenesis kit (NEB) and appropriate primer sets (**Table S4**). All plasmid sequences were verified by Sanger sequencing (ElimBio).

### *E. coli* Growth Assay

NEB Express I^q^ cells were transformed with each modified pLK35 plasmid with the desired 23S rRNA mutations. Liquid cultures of each transformant were grown overnight, and the resulting cultures were diluted to OD 0.01 in LB media containing 100 µg/mL ampicillin and 500 µM IPTG. 200 µL of each dilution was then pipetted into wells of a sterile 96-well plate. The OD_600_ was measured every 15 minutes using a Spark Plate Reader (Tecan) at 37 °C with constant shaking. Doubling times were calculated by fitting the initial region of exponential growth to a Malthusian growth model in Prism.

To determine the ratio of tagged ribosomes to WT ribosomes present during the growth experiment, 200 µL of culture from each sample was pelleted. Total RNA was extracted from cell pellets using the RNeasy Mini Kit (Qiagen). Semiquantitative PCR was used to determine the ratio of tagged 23S rRNA to WT 23S rRNA as previously described.^38^ Briefly, primer MS2_quant_R was used to reverse transcribe a portion of 23S rRNA. A region containing the sequence for the MS2 tag was amplified via PCR using primers MS2_quant_F and MS2_quant_R. PCR products were run on a 10% polyacrylamide-TBE gel (Invitrogen) and visualized with SYBR gold stain (ThermoFisher). DNA bands were quantified using ImageJ software.^46^

### Crude Ribosome Preparation

Crude ribosomes were isolated as previously described with several adaptations.^47^ NEB Express I^q^ cells were transformed with the corresponding pLK35 plasmid carrying a tac promoter, MS2-tagged 23S rRNA, and the desired mutations. Overnight cultures of the transformants were diluted 1:100 into 2-3 L of LB broth containing 100 µg/mL ampicillin and grown at 37 °C. Upon growth to OD_600_=0.6, rRNA transcription was induced with 0.5 mM IPTG and the cultures were grown for 3 hours at 37 °C. Cells were then pelleted, resuspended in 50-100 mL buffer A (20 mM Tris-HCl pH 7.5, 100 mM NH_4_Cl, 10 mM MgCl_2_, 0.5 mM EDTA, 2 mM DTT), and lysed by sonication. The lysate was then clarified by centrifugation at 14,000 rpm (34,000 xg) for 45 minutes in a F14-14×50cy rotor (ThermoFisher). Clarified lysate was then loaded onto a sucrose cushion containing 24 mL of buffer B (20 mM Tris-HCl pH 7.5, 500 mM NH_4_Cl, 10 mM MgCl_2_, 0.5 mM EDTA, 2 mM DTT) with 0.5 M sucrose and 17 ml of buffer C (20 mM Tris-HCl pH 7.5, 60 mM NH_4_Cl, 6 mM MgCl_2_, 0.5 mM EDTA, 2 mM DTT) with 0.7 M sucrose in Ti-45 tubes (Beckman-Coulter). The tubes were then placed in a Ti-45 rotor and spun at 27,000 rpm (57,000 xg) for 16 hours at 4 °C. The next day, ribosome pellets were resuspended in dissociation buffer (buffer C with 1 mM MgCl_2_).

### MS2-tagged Ribosome Purification

The MBP-MS2 fusion protein was purified as previously described.^38^ 10 mg of MBP-MS2 protein was diluted to 1 mg/mL in MS2-150 buffer (20 mM HEPES pH 7.5, 150 mM KCl, 1 mM EDTA, 2 mM 2-mercaptoethanol) for a 5 mL column preparation. For each 1 mL column prep, 4 mg of MBP-MS2 protein was used. The 5 mL or 1 mL MBP Trap column (Cytiva) was washed with 10 column volumes (CV) of MS2-150 buffer and then the MBP-MS2 protein was loaded slowly onto the column. The column was then washed with 5 CV of buffer A-1 (buffer A with 1 mM MgCl_2_). Crude tagged ribosomes (∼100-200 mg) were diluted to 15 mg/mL in buffer A-1, which dissociates 70S ribosomes into 30S and 50S subunits and decreases WT ribosome contamination. The diluted crude ribosomes were loaded slowly onto the MBP-trap column, which was then washed with 5 CV of buffer A-1 followed by 5 CV of buffer A-250 (buffer A with 250 mM NH_4_Cl and 1 mM MgCl_2_). Ribosomes were eluted with a 10 CV gradient of buffer A-1 containing 0-10 mM maltose. Fractions containing tagged ribosomes were concentrated in 100 kDa cut off spin filters (Millipore) and washed with buffer A-1. 50S ribosomal subunits were quantified using the approximation of 1 A_260_= 36 nM and were stored at -80 °C.

### Untagged Subunit Purification

Overnight cultures of *E. coli* strain MRE600 were diluted 1:100 into 2-3 L of LB broth and grown to an OD_600_=0.6. Cells were then pelleted and lysed by sonication. Ribosomes were pelleted and resuspended as described above for crude ribosome preparations. After resuspension in dissociation buffer, the 30S and 50S subunits were separated on a 15-35% (w/v) sucrose gradient in dissociation buffer on a SW-32 rotor (Beckman-Coulter) spun at 28,000 rpm (97,000 xg) for 16 hours. Fractions containing the 50S and 30S subunits were collected separately, washed with dissociation buffer, and concentrated in a 100 kDa cut off spin filter. To increase purity, concentrated subunits were separately run on second 15-35% sucrose gradients and appropriate fractions were washed with dissociation buffer and concentrated.

### Nanoluciferase 11S Fragment Purification

A plasmid encoding 8xHis-tagged 11S was transformed into BL21 (DE3) Rosetta2 pLysS cells (Macrolab, UC Berkeley). An overnight culture was diluted (1:100) into LB media with 100 µg/mL ampicillin and grown at 37 °C. When the culture reached OD_600_=0.4, cells were induced with 1 mM IPTG for three hours. Cells were then pelleted at 4,000 xg for 15 minutes and resuspended in ∼20 mL 11S lysis buffer (20 mM HEPES pH 7.5, 50 mM KCl, 10% glycerol, and 10 mM imidazole). Cells were lysed by sonication and the lysate was clarified by centrifugation at 18,000 rpm (25,000 xg) for 30 minutes (JA-20 rotor, Beckman). Supernatant was then applied to a 1 mL HisTrap column (Cytiva). The column was washed with 10 CV of 11S lysis buffer and eluted with a 20 CV linear gradient form 20-500 mM imidazole in lysis buffer. Protein fractions were dialyzed against 11S lysis buffer without imidazole overnight at 4 °C. The protein was then concentrated and stored at -80 °C.

### HiBit Translation Assay

50S ribosomal subunits were diluted to 1.4 µM in buffer A with a final concentration of 10 mM MgCl_2_. This mixture was then incubated at the desired temperature for 30 minutes on a ProFlex (ThermoFisher) PCR cycler with a heated lid. After incubation, tubes containing the 50S subunits were placed on the bench and allowed to cool slowly to room temperature for 30 minutes. The cooling method influenced the return of mutant ribosome activity. Slow cooling at room temperature returned higher levels of activity as compared to snap-cooling on ice for the CC mutant ribosome (**Figure S12**). It is possible that snap-cooling leads to a less active conformation in the mutant ribosome. Due to this observation, a slow-cooling method was utilized. After cooling, an *in vitro* translation mixture was assembled using a ∆Ribosome PURExpress kit (NEB). The mixture was assembled with the following: 3.2 µL solution A (NEB), 1 µL factor mix (NEB), 250 nM pre-incubated 50S ribosomal subunits, 500 nM WT untagged 30S ribosomal subunits, 1U/µL Murine RNAse inhibitor (NEB), 100 nM 11S NanoLuc protein^23^, 1:50 (v/v) dilution of Nano-Glo substrate (Promega), and 10 ng/µL of DNA template containing a T7 promoter, ribosome binding site, and the coding sequence for the HiBit peptide^23^ (final volume of 8 µL). 2 µL of the *in-vitro* translation mixture was placed in a 384 well plate, and luminescence was measured as a function of time in a Spark Plate Reader (Tecan). Ribosome activity was calculated by determining the slope of the initial linear region of each *in vitro* translation reaction.

### *In Vitro* nLuc Translation Endpoint Assay

An *in vitro* translation reaction was assembled with the following: 6.4 µL solution A (NEB), 2 µL factor mix (NEB), 250 nM pre-incubated 50S subunit, 500 nM WT untagged 30S subunit, 1U/µL Murine RNAse inhibitor (NEB), and 10 ng/µL of a plasmid encoding a T7 promoter followed by the nLuc gene (final volume of 16 µL). This reaction was then incubated at 37 °C for 1 hour. After incubation, 5 µL from the reaction was mixed with 30 µL of 11S buffer (20 mM HEPES pH 7.5, 50 mM KCl, and 10% glycerol) and a 1:50 dilution of Nano-Glo substrate (Promega). This mixture was placed in a 384 well plate and luminescence was measured in a Spark Plate Reader (Tecan).

### Sucrose Gradients

For heat treated 50S subunits, 10 pmol 50S subunit was diluted in buffer A with 10 mM MgCl_2_ to a final volume of 20 µL. This mixture was then incubated at the desired temperature for 30 minutes on a ProFlex (ThermoFisher) PCR cycler with a heated lid and placed on the bench for 30 minutes to cool slowly to room temperature. For 70S formation assays, 10 pmol 50S subunit and 20 pmol WT untagged 30S subunit were diluted in buffer A with 15 mM MgCl_2_ to a final volume of 20 µL. These samples were then incubated at 37 °C for 30 minutes. Samples were loaded onto a 15-40% sucrose gradient in buffer A with 10 or 15 mM MgCl_2_. Gradients were then placed in a SW-41 Ti rotor (Beckman-Coulter) and centrifuged at 29,000 rpm (100,000 xg) for 13 hours. An ISCO gradient fractionation system was used to measure A_254_ traces.

### CD Spectroscopy

RNA oligonucleotides were purchased from IDT (**Table S5**) and resuspended into RNAse-free water. RNA was incubated at 95 °C for 3 minutes and then immediately snap-cooled in an ice water bath. After cooling, the RNA concentration was adjusted to 100 µM and the buffer concentration was adjusted to 20 mM K_x_H_y_PO_4_ pH 6.5. Low ionic buffer conditions were used to shift the melting temperature to a value where both the initial and final baselines could be more accurately modeled. CD experiments were conducted in a 1-mm path-length cuvette with an Aviv 410 CD spectropolarimeter. For thermal melting experiments, CD at 262 nm was recorded in 2 °C intervals from 5 °C to 95 °C. The temperature was moved to the target value, and the system was allowed to equilibrate for one minute before a measurement was taken. Thermal melting data was fit to a two-state model with sloping baselines^48^ (**Equation 1**) in Prism 9. All RNAs tested exhibited reversible thermal melting profiles.

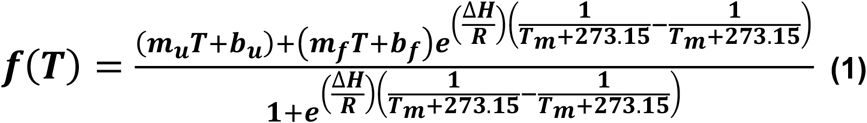

Where T is temperature in Celsius, m_u_ and m_f_ are the slopes of the unfolded and folded baselines, respectively, b_u_ and b_f_ are the y-intercepts of the unfolded and folded baselines, respectively, ∆H is the Vant Hoff enthalpy of unfolding, R is the gas constant, and T_m_ is the melting temperature in Celsius.

### Methionyl-tRNA Synthetase Purification

A plasmid encoding 6xHis-tagged methionyl-tRNA synthetase (MetRS) was transformed into BL21 (DE3) Codon+ RIL cells (Agilent). Overnight cultures were diluted (1:100) into ZYM-5052 autoinducing media^49^ and grown overnight at 37 °C. Cells were pelleted at 4,000 xg for 15 minutes and resuspended in 50 mL of MetRS lysis buffer (20 mM Tris pH 7.5, 150 mM NaCl, 5 mM imidazole, and 0.5 mM EDTA). Cells were lysed by sonication and the lysate was clarified by centrifugation at 18,000 rpm (25,000 xg JA-20 rotor, Beckman-Coulter) for 30 minutes. The supernatant was applied to a 5 mL HisTrap column (Cytiva), washed with 5 CV of MetRS lysis buffer with 23 mM imidazole and eluted with a 20 CV linear gradient from 23-500 mM imidazole. Protein fractions were pooled and dialyzed overnight against MetRS buffer (50 mM HEPES pH 7.5, 100 mM KCl, 10 mM MgCl_2_, 7 mM BME and 30% glycerol). The protein was concentrated and stored at -80 °C.

### tRNA Preparation

A single stranded DNA template encoding a T7 promoter and tRNA^fMet^, with a C1G mutation to increase transcription yield and deletion of the 5’
s-terminal adenosine (C1G -A), was amplified via PCR, yielding a double stranded DNA template. tRNA^fMet^ was *in vitro* transcribed with transcription buffer (50 mM Tris pH 7.5, 15 mM MgCl_2_, 5 mM DTT, 2 mM spermidine), 2.5 mM NTPs, 1 U/µL Murine RNase inhibitor (NEB), 0.5 U/µL T7 polymerase (NEB), and 0.5 U/mL YIPP (NEB). *In vitro* transcription reactions were incubated for 16 hours at 37 °C and then treated with 45 U/mL of RQ1 DNase (Promega) at 37 °C for 30 minutes. Reactions were ethanol precipitated and then resolved on a 12% denaturing polyacrylamide gel. Gel slices containing the tRNA were excised, crushed, and soaked in 300 mM sodium acetate pH 5.2 overnight at 4 °C. The RNA was then ethanol precipitated, resuspended in water, and stored at -80 °C.

tRNA^fMet^ lacking the 3’
s-terminal adenosine (A76) was then treated with the tRNA nucleotidyl transferase from *Archeoglobus fulgidus* (*Af*) and 3′-NH_2_-ATP (nATP) (Axxora) in the following conditions: 100 mM glycine pH 9.0, 10 mM MgCl_2_, 2 µM nucleotidyl transferase, 2 µM tRNA(-A), 0.5 mM nATP, 1 mM DTT, and 2 U/mL YIPP (NEB).^50^ *Af* tRNA nucleotidyl transferase was purified with the same method and buffers as the MBP-MS2 fusion protein.^38^ The reaction was incubated at 37 °C for 2 hours. The tRNA was then phenol-chloroform extracted, ethanol precipitated, and resuspended in water.

*E. coli* methionyl-tRNA synthetase (MetRS) was used to charge 3’
s-amino tRNA^fMet^ in the following conditions: charging buffer (50 mM HEPES pH 7.5, 10 mM KCl, 20 mM MgCl_2_, 2 mM DTT), 10 µM tRNA^fMet^-NH_2_, 10 mM methionine, 1 µM MetRS, 10 mM ATP, 1 U/µL Murine RNase inhibitor. The reaction was incubated at 37 °C for 30 minutes. Charged tRNA was phenol-chloroform extracted, ethanol precipitated, resuspended in water, and stored at -80 °C.

### Cryo-EM Sample Preparation

To assemble 70S ribosomes, 0.5 mg of U2554C U2555C (CC) mutant 50S subunit and 0.6 mg of WT untagged 30S subunit were incubated in buffer C with 10 mM MgCl_2_ at 37 °C for 45 minutes. The ribosome mixture was then loaded onto a 15-40% (w/v) sucrose gradient in buffer C with 10 mM MgCl_2_. Gradients were centrifuged at 28,000 rpm (97,000 xg) for 16 hours in a SW-32 rotor (Beckman-Coulter). An ISCO gradient fraction system was used to isolate the 70S fraction (**Figure S13**).

Ribosome-tRNA-mRNA complexes were prepared as previously described^39^ with the following change. The mRNA used had the sequence 5’
s-GUAUAAGGAGGUAAA<u>AUGAUG</u>UAACUA-3′ (IDT). Met codons are underlined. Paromomycin at a concentration of 100 µM was used to ensure nonenzymatic A-site tRNA binding. Cryo-EM grids were prepared and samples were frozen as previously described.^39^

### Cryo-EM Data Acquisition, Image Processing, and Modeling

Movies were collected on a 300 kV Titan Krios microscope with a GIF energy filter and a Gatan K3 camera, and data was acquired as described.^39^ Raw movies were binned to the physical pixel size (0.8293 Å) in RELION 3.1^51^ and motion corrected with MotionCor2^52^. The CTFs of micrographs were estimated using CTFFind4^53^. The estimated CTFs of micrographs were visually inspected, and micrographs with poor CTF fit were manually rejected. Particles were picked using the RELION implementation with a Laplacian of Gaussian method, and then extracted and binned 4x for classification steps. 2D classification was performed in RELION using 150 classes. Junk particles were rejected and a second round of 2D classification was performed. 3D classification was performed with a cryoSPARC heterogeneous refinement^54^ with 10 classes using an initial model generated in EMAN2^55^ from PDB 1VY4^20^. Classes containing features consistent with well-ordered 70S particles were combined and a local refinement on the 50S subunit was carried out. The aligned particles were then transferred back to RELION. A 3D classification without alignment was performed in RELION to separate classes with slightly rotated 30S subunits. This procedure yielded a junk class, a rotated 30S subunit that had no A-site tRNA density, and a class that had P-site and A-site tRNA density. The latter class was then subjected to an additional classification without alignment using a mask on the A-site tRNA. Interestingly, this step yielded two slightly shifted A-site tRNA classes that were equally populated (**Figure S3**). The separation of multiple A-site tRNA classes with a masked 3D classification has been observed with the WT ribosome.^21^ Upon closer examination, one of these classes had clear density for the CCA-end of the A-site tRNA and was processed further. These particles were re-extracted at full size and subjected to CTF Refinement^51^ and Bayesian polishing^56^ in RELION. Finally, a focus refinement on the 50S subunit was performed to generate the final map. PDB 7K00 was used as an initial 70S ribosome model^39^ and the ‘Fit to Map’ function in ChimeraX^57^ was used to align the cryo-EM map to the initial model. The 70S model was refined using real space refinement in PHENIX.^58^ The model and cryo-EM map were inspected in COOT^59^ and manual adjustments to the model were made as needed.

## Supporting information

PDB coordinates

cryo-EM map fragment

## Data Availability

Sequences, temperature data, and scripts used in the sequence and phylogenetic analysis are available at https://github.com/petaripenev/A-loop_sequence_analysis.

## Acknowledgments

We thank Dan Toso and Paul Tobias for help with cryo-EM data collection, Susan Marqusee and her lab for the use of their CD spectrometer, and Fred Ward for help with protein purification. This work was funded primarily by the NSF Center for Genetically Encoded Materials (C-GEM), CHE-2002182. AJN was partially supported by an NIH training grant (T32 066698). PIP and JFB were supported by the Innovative Genomics Institute at Berkeley and the Chan Zuckerberg Biohub.

## REFERENCES

1. Porse, B. T. & Garrett, R. A. Mapping important nucleotides in the peptidyl transferase centre of 23 S rRNA using a random mutagenesis approach. J. Mol. Biol. 249, 1–10 (1995).

2. Gregory, S. T., Lieberman, K. R. & Dahlberg, A. E. Mutations in the peptidyl transferase region of E.coli 23s rRNA affecting translational accuracy. Nucleic Acids Res. 22, 279–284 (1994).

3. Thompson, J. et al. Analysis of mutations at residues A2451 and G2447 of 23S rRNA in the peptidyltransferase active site of the 50S ribosomal subunit. Proc. Natl. Acad. Sci. U. S. A. 98, 9002–9007 (2001).

4. D’Aquino, A. E. et al. Mutational characterization and mapping of the 70S ribosome active site. Nucleic Acids Res. 48, 2777–2789 (2020).

5. Davidovich, C., Bashan, A. & Yonath, A. Structural basis for cross-resistance to ribosomal PTC antibiotics. Proc. Natl. Acad. Sci. U. S. A. 105, 20665–20670 (2008).

6. Steitz, T. A. A structural understanding of the dynamic ribosome machine. Nat. Rev. Mol. Cell Biol. 9, 242–253 (2008).

7. Kim, D. F. & Green, R. Base-Pairing betwen 23S rRNA and tRNA in the RIbosomal A Site. Mol. Cell 4, 859–864 (1999).

8. Terasaka, N., Hayashi, G., Katoh, T. & Suga, H. An orthogonal ribosome-tRNA pair via engineering of the peptidyl transferase center. Nat. Chem. Biol. 10, 555–557 (2014).

9. Bernier, C. R., Petrov, A. S., Kovacs, N. A., Penev, P. I. & Williams, L. D. Translation: The universal structural core of life. Mol. Biol. Evol. 35, 2065–2076 (2018).

10. Wang, W. et al. Loss of a single methylation in 23s rRNA delays 50s assembly at multiple late stages and impairs translation initiation and elongation. PNAS 117, 15609–15619 (2020).

11. Caldas, T., Binet, E., Bouloc, P. & Richarme, G. Translational Defects of Escherichia coli Mutants Deficient in the Um2552 23S Ribosomal RNA Methyltransferase RrmJ/FTSJ. Biochem. Biophys. Res. Commun. 271, 714–718 (2000).

12. Bashan, A. et al. Structural basis of the ribosomal machinery for peptide bond formation, translocation, and nascent chain progression. Mol. Cell 11, 91–102 (2003).

13. O’Connor, M. & Dahlberg, A. E. Mutations at U2555, a tRNA-protected base in 23S rRNA, affect translational fidelity. PNAS 90, 9214–9218 (1993).

14. Jegousse, C., Yang, Y., Zhan, J., Wang, J. & Zhou, Y. Structural signatures of thermal adaptation of bacterial ribosomal RNA, transfer RNA, and messenger RNA. PLoS One 12, 1–14 (2017).

15. Sas-Chen, A. et al. Dynamic RNA acetylation revealed by quantitative cross-evolutionary mapping. Nature 583, 638–643 (2020).

16. Noon, K. R., Bruenger, E. & McCloskey, J. A. Posttranscriptional modifications in 16S and 23S rRNAs of the archaeal hyperthermophile Sulfolobus solfataricus. J. Bacteriol. 180, 2883–2888 (1998).

17. Quast, C. et al. The SILVA ribosomal RNA gene database project: Improved data processing and web-based tools. Nucleic Acids Res. 41, 590–596 (2013).

18. Brochier-Armanet, C., Boussau, B., Gribaldo, S. & Forterre, P. Mesophilic crenarchaeota: Proposal for a third archaeal phylum, the Thaumarchaeota. Nat. Rev. Microbiol. 6, 245–252 (2008).

19. Youngman, E. M. & Green, R. Affinity purification of in vivo-assembled ribosomes for in vitro biochemical analysis. Methods 36, 305–312 (2005).

20. Polikanov, Y. S., Steitz, T. A. & Innis, C. A. A proton wire to couple aminoacyl-tRNA accommodation and peptide-bond formation on the ribosome. Nat. Struct. Mol. Biol. 21, 787–793 (2014).

21. Watson, Z. L. et al. All-atom metadynamics captures preferred geometries of non-L-a-amino acid monomers within the ribosomal PTC. In preparation (2022).

22. Sokoloski, J. E., Godfrey, S. A., Dombrowski, S. E. & Bevilacqua, P. C. Prevalence of syn nucleobases in the active sites of functional RNAs. Rna 17, 1775–1787 (2011).

23. Dixon, A. S. et al. NanoLuc Complementation Reporter Optimized for Accurate Measurement of Protein Interactions in Cells. ACS Chem. Biol. 11, 400–408 (2016).

24. Thompson, J. & Dahlberg, A. E. Testing the conversation of the translational machinery over evolution in diverse environments: Assaying Thermus thermophilus ribosomes and initiation factors in a coupled transcription-translation system from Escherichia coli. Nucleic Acids Res. 32, 5954–5961 (2004).

25. Metelev, M. et al. Direct measurements of mRNA translation kinetics in living cells. Nat. Commun. 13, 1–13 (2022).

26. Lancaster, L., Lambert, N. J., Maklan, E. J., Horan, L. H. & Noller, H. F. The sarcin-ricin loop of 23S rRNA is essential for assembly of the functional core of the 50S ribosomal subunit. Rna 14, 1999–2012 (2008).

27. Dedkova, L. M., Fahmi, N. E., Golovine, S. Y. & Hecht, S. M. Enhanced D-amino acid incorporation into protein by modified ribosomes. J. Am. Chem. Soc. 125, 6616–6617 (2003).

28. Melo Czekster, C., Robertson, W. E., Walker, A. S., Söll, D. & Schepartz, A. In Vivo Biosynthesis of a β-Amino Acid-Containing Protein. J. Am. Chem. Soc. 138, 5194–5197 (2016).

29. Youngman, E. M., Brunelle, J. L., Kochaniak, A. B. & Green, R. The active site of the ribosome is composed of two layers of conserved nucleotides with distinct roles in peptide bond formation and peptide release. Cell 117, 589–599 (2004).

30. Ohira, T. et al. Reversible RNA phosphorylation stabilizes tRNA for cellular thermotolerance. Nature 605, 372–379 (2022).

31. Rakauskaite, R. & Dinman, J. D. rRNA mutants in the yeast peptidyltransferase center reveal allosteric information networks and mechanisms of drug resistance. Nucleic Acids Res. 36, 1497–1507 (2008).

32. O’Connor, M., Willis, N. M., Bossi, L., Gesteland, R. F. & Atkins, J. F. Functional tRNAs with altered 3’ ends. EMBO J. 12, 2559–2566 (1993).

33. Blanchard, S. C. & Puglisi, J. D. Solution structure of the A loop of 23S ribosomal RNA. PNAS 98, 3720–3725 (2001).

34. Nikolay, R. et al. Structural Visualization of the Formation and Activation of the 50S Ribosomal Subunit during In Vitro Reconstitution. Mol. Cell 70, 881-893.e3 (2018).

35. Bloom, J. D., Labthavikul, S. T., Otey, C. R. & Arnold, F. H. Protein stability promotes evolvability. Proc. Natl. Acad. Sci. U. S. A. 103, 5869–5874 (2006).

36. Grasso, K. T. et al. Structural Robustness Affects the Engineerability of Aminoacyl-tRNA Synthetases for Genetic Code Expansion. Biochemistry 60, 489–493 (2021).

37. Kofman, C., Watkins, A. M., Soon, D. & Alexandra, C. Computationally-guided design and selection of ribosomal active site mutants with high activity. bioRxiv 1–17 (2022).

38. Ward, F. R., Watson, Z. L., Ad, O., Schepartz, A. & Cate, J. H. D. Defects in the Assembly of Ribosomes Selected for β-Amino Acid Incorporation. Biochemistry 58, 4494–4504 (2019).

39. Watson, Z. L. et al. Structure of the bacterial ribosome at 2 Å resolution. Elife 9, 1–62 (2020).

40. Katoh, K., Misawa, K., Kuma, K. I. & Miyata, T. MAFFT: A novel method for rapid multiple sequence alignment based on fast Fourier transform. Nucleic Acids Res. 30, 3059–3066 (2002).

41. Katoh, K., Rozewicki, J. & Yamada, K. D. MAFFT online service: Multiple sequence alignment, interactive sequence choice and visualization. Brief. Bioinform. 20, 1160–1166 (2018).

42. Sato, Y., Okano, K., Kimura, H. & Honda, K. Tempura: Database of growth temperatures of usual and rare prokaryotes. Microbes Environ. 35, 1–3 (2020).

43. Letunic, I. & Bork, P. Interactive tree of life (iTOL) v5: An online tool for phylogenetic tree display and annotation. Nucleic Acids Res. 49, W293–W296 (2021).

44. Rinke, C. et al. A standardized archaeal taxonomy for the Genome Taxonomy Database. Nat. Microbiol. 6, 946–959 (2021).

45. Douthwaite, S., Powers, T., Lee, J. Y. & Noller, H. F. Defining the Structural Requirements for a Helix in 23S Ribosomal RNA that Confers Erythromycin Resistance. J. Mol. Biol. 209, 655–665 (1989).

46. Schneider, C. A., Rasband, W. S. & Eliceiri, K. W. NIH Image to ImageJ: 25 years of image analysis. Nat. Methods 9, 671–675 (2012).

47. Travin, D. Y. et al. Structure of ribosome-bound azole-modified peptide phazolicin rationalizes its species-specific mode of bacterial translation inhibition. Nat. Commun. 10, (2019).

48. Kotar, A., Ma, S. & Keane, S. C. pH dependence of C•A, G•A and A•A mismatches in the stem of precursor microRNA-31. Biophys. Chem. 283, (2022).

49. Studier, F. W. Stable expression clones and auto-induction for protein production in E. coli. Methonds Mol. Biol. 1091, 17–32 (2014).

50. Katoh, T. & Suga, H. Flexizyme-catalyzed synthesis of 3′-aminoacyl-NH-tRNAs. Nucleic Acids Res. 47, e54 (2019).

51. Zivanov, J. et al. New tools for automated high-resolution cryo-EM structure determination in RELION-3. Elife 7, 1–22 (2018).

52. Zheng, S. Q. et al. MotionCor2: anisotropic correction of beam-induced motion for improved cryo-electron microscopy. Nat. Methods 14, 331–332 (2017).

53. Rohou, A. & Grigorieff, N. CTFFIND4: Fast and accurate defocus estimation from electron micrographs. J. Struct. Biol. 192, 216–221 (2015).

54. Punjani, A., Rubinstein, J. L., Fleet, D. J. & Brubaker, M. A. cryoSPARC: algorithms for rapid unsupervised cryo-EM structure determination. Nat. Methods 14, 290–296 (2017).

55. Tang, G. et al. EMAN2: An extensible image processing suite for electron microscopy. J. Struct. Biol. 157, 38–46 (2007).

56. Zivanov, J., Nakane, T. & Scheres, S. H. W. A Bayesian approach to beam-induced motion correction in cryo-EM single-particle analysis. IUCrJ 6, 5–17 (2019).

57. Pettersen, E. F. et al. UCSF ChimeraX: Structure visualization for researchers, educators, and developers. Protein Sci. 30, 70–82 (2021).

58. Liebschner, D. et al. Macromolecular structure determination using X-rays, neutrons and electrons: Recent developments in Phenix. Acta Crystallogr. Sect. D Struct. Biol. 75, 861–877 (2019).

59. Casañal, A., Lohkamp, B. & Emsley, P. Current developments in Coot for macromolecular model building of Electron Cryo-microscopy and Crystallographic Data. Protein Sci. 29, 1069–1078 (2020).

