## Supplemental Information for "Rare Ribosomal RNA Sequences from Archaea Stabilize the Bacterial Ribosome"

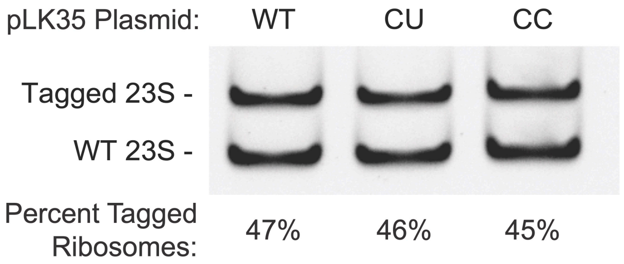


**Figure S1. Percentage of MS2-tagged ribosomes in *E. coli* expressing 23S rRNA variants.** Following *E. coli* growth assays, 200 μL of induced culture was isolated to detect the ratio of MS2-tagged ribosomes to WT endogenous ribosomes at the end of the experiment. RT-PCR samples were run on a 10% polyacrylamide-TBE gel. DNA containing an MS2 tag sequence is 32 bp larger than DNA with the native 23S sequence. Amplified bands were quantified to determine the percent of MS2-tagged ribosomes.

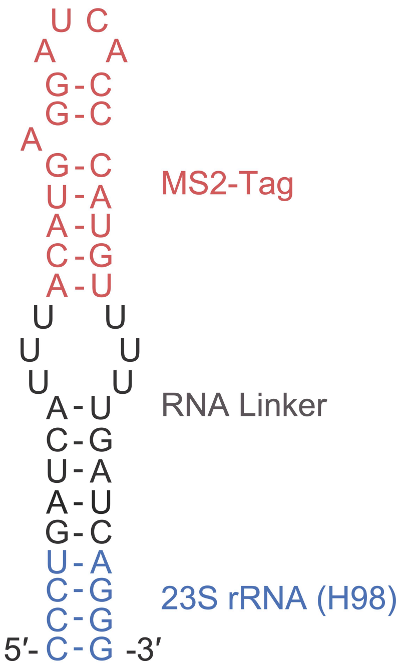


**Figure S2. MS2-Tag design used in this study.** The MS2 tag is inserted into rRNA helix H98 in the 50S *E. coli* ribosome. The H98 stem is elongated and the MS2 stem loop is attached with a poly-U linker.

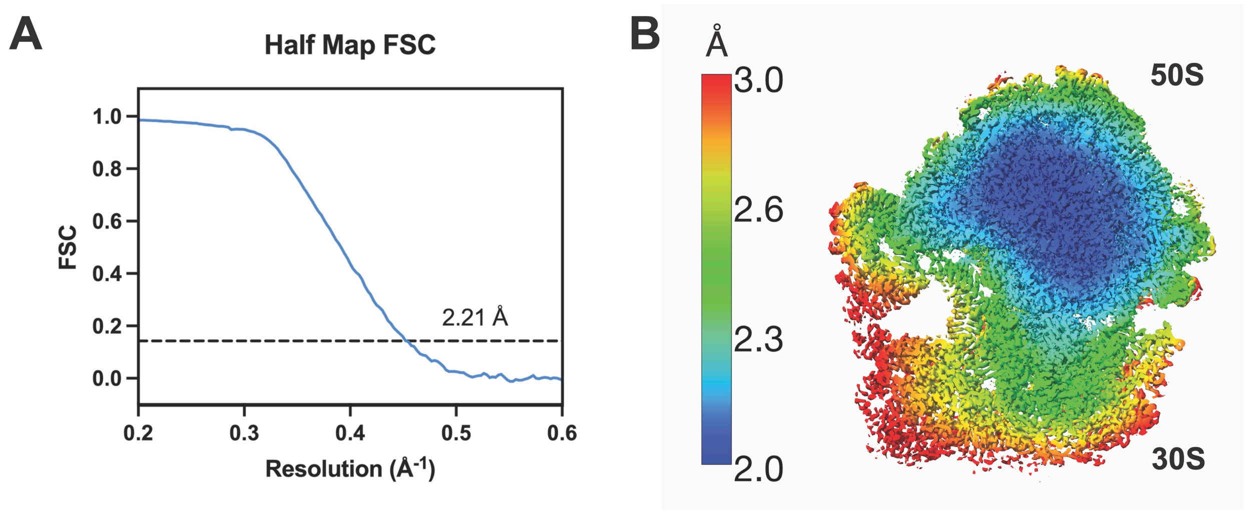


**Figure S3. Resolution of the cryo-EM map generated from the particles in Class I.** A) The global FSC resolution of the CC 70S ribosome is 2.2 Å. B) Local resolution is plotted on the 70S ribosome map.

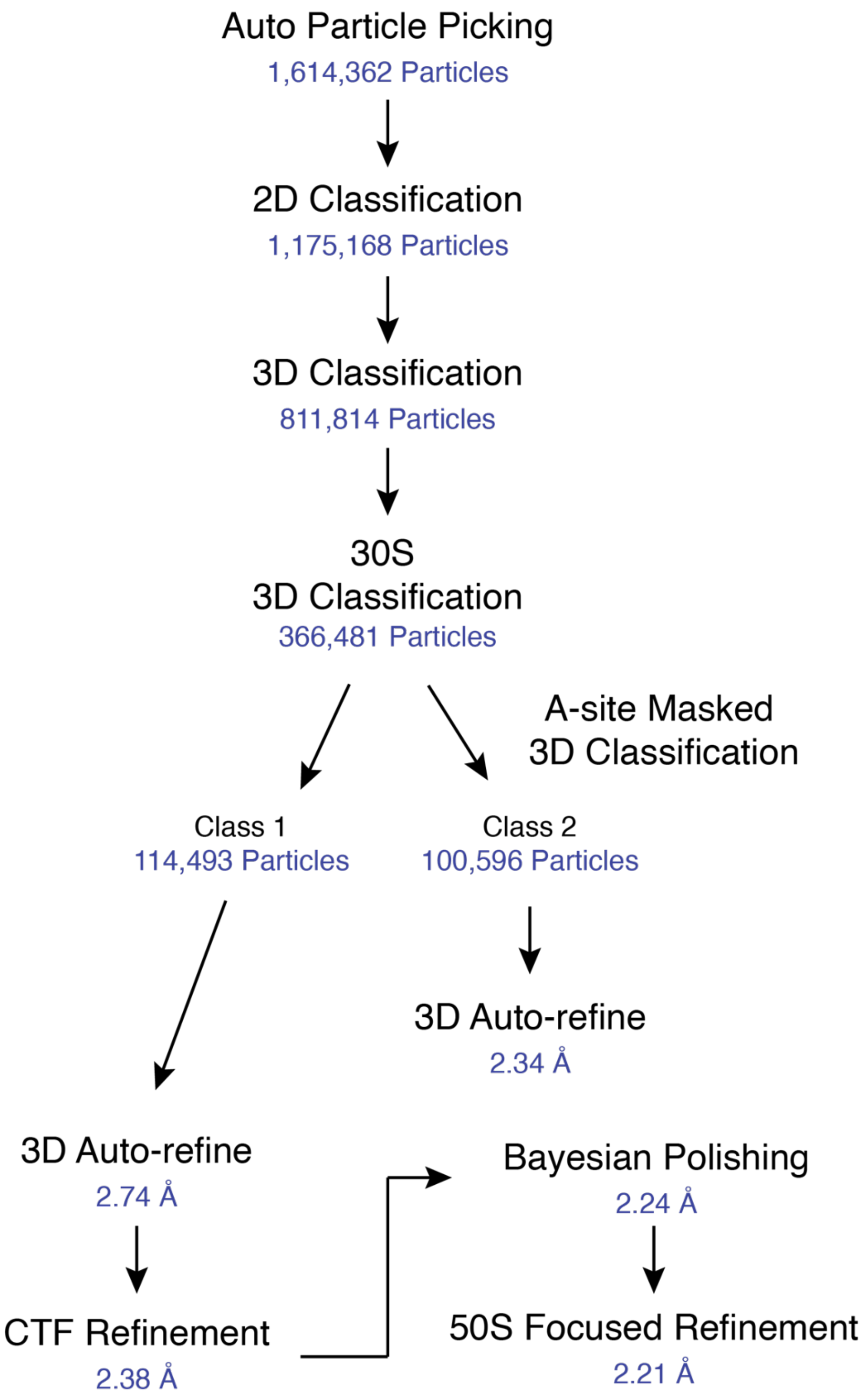


**Figure S4. Cryo-EM processing workflow.** A-site Class 1 had density for the tRNA 3'-CCA end and was further refined. Class 2 lacked density for the A-site tRNA 3´-CCA end.

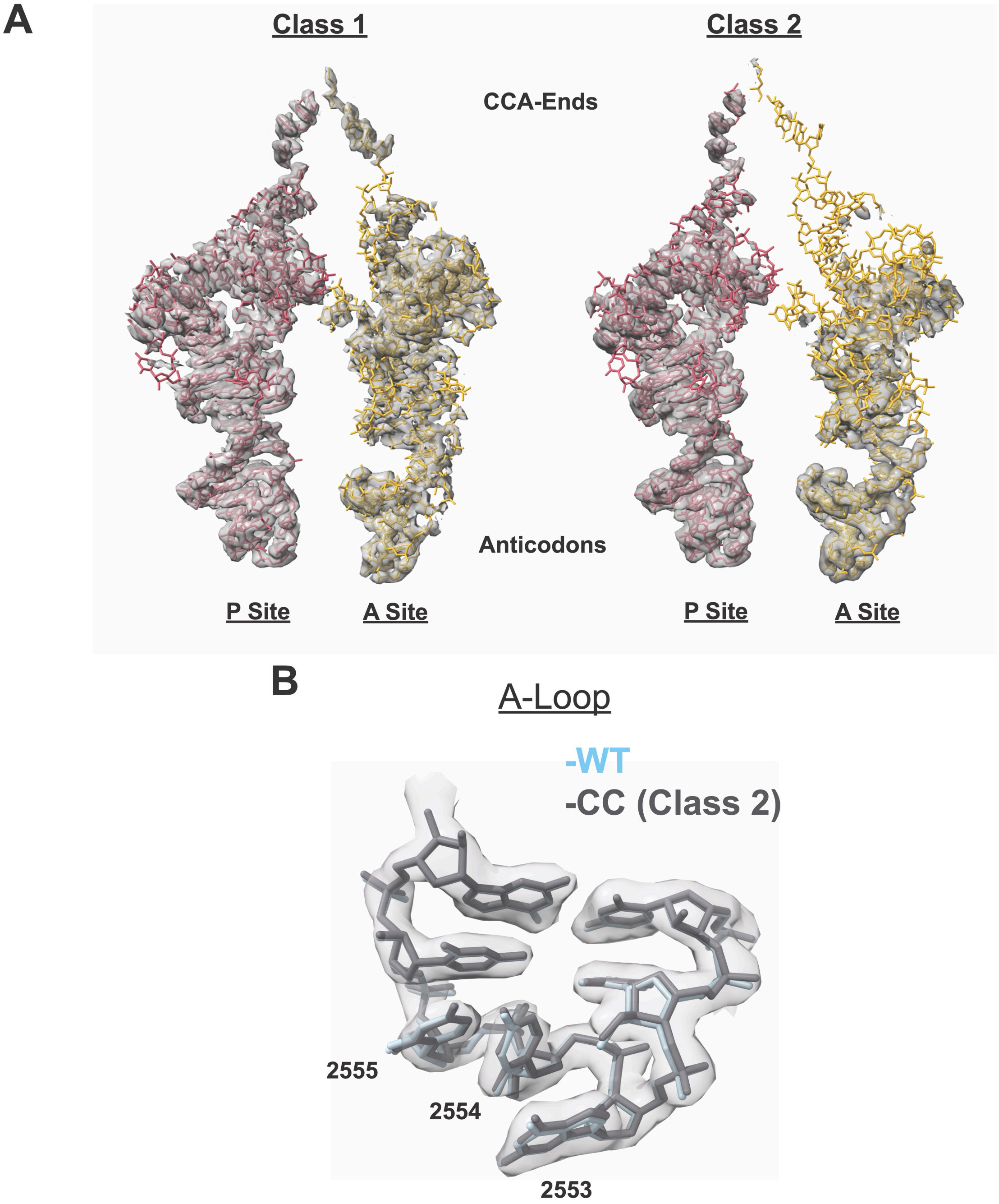


**Figure S5. Features of the tRNAs in A-site classes 1 and 2.** A) P-site and A-site tRNA model and cryo-EM density for A-site class 1 (left) and class 2 (right). Class 2 corresponded to an A-site tRNA class that lacked density for most of the acceptor stem. A B-factor of 30 Å$^2$ was applied to the cryo-EM maps. B) Cryo-EM density and model for the A-loop from Class 2 are shown in grey. The model from a WT ribosome structure[1] is shown in light blue.

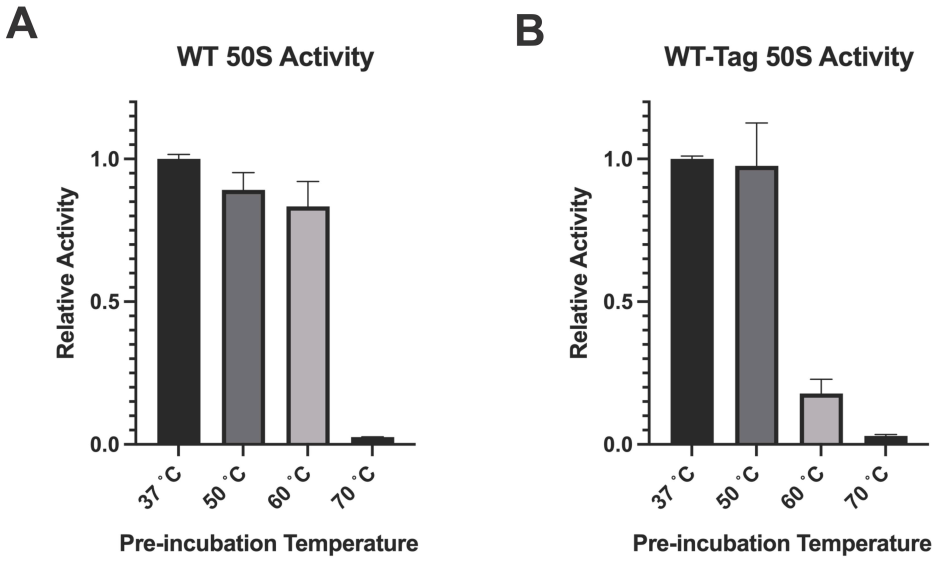


**Figure S6. Activity of heat-treated 50S subunits without and with the MS2 tag.** A) Untagged and B) MS2-tagged WT 50S ribosomes were pre-incubated at the indicated temperatures for 30 minutes and then slow cooled for 30 minutes. The 50S ribosomes were then used in the HiBit translation assay.

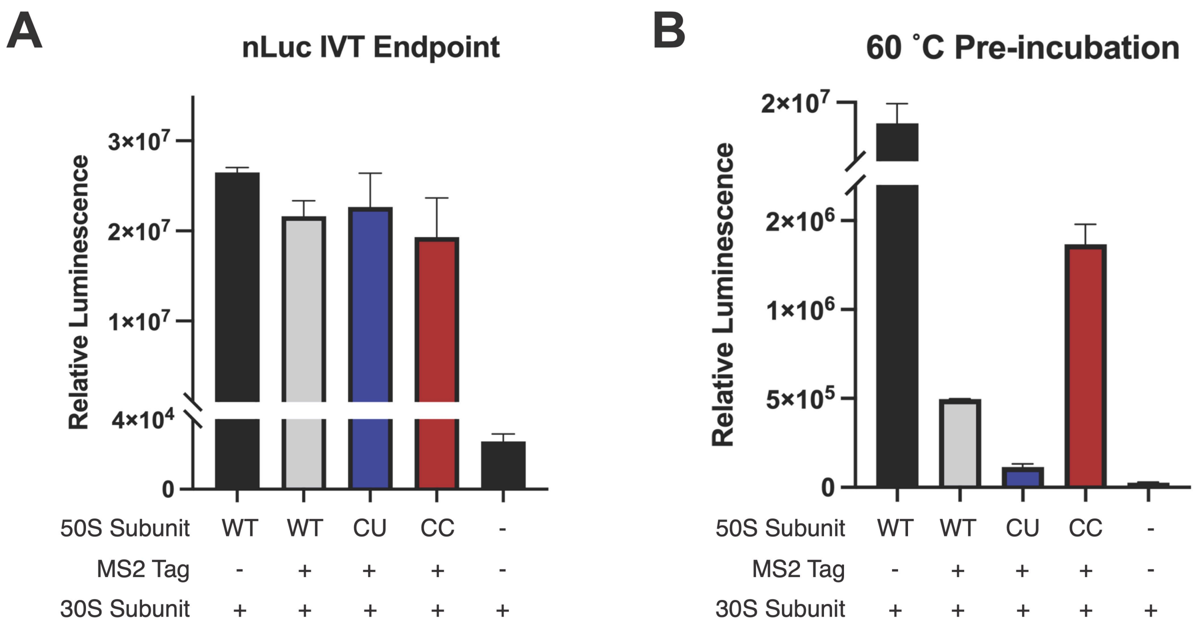


**Figure S7. Activity of ribosomes in nLuc translation endpoint assays.** A) Replicate of an nLuc IVT endpoint assay at 37 °C for CU and CC ribosomes. B) 50S ribosomes were preincubated at 60 °C and cooled slowly. Their relative activities were then determined with the nLuc endpoint assay.

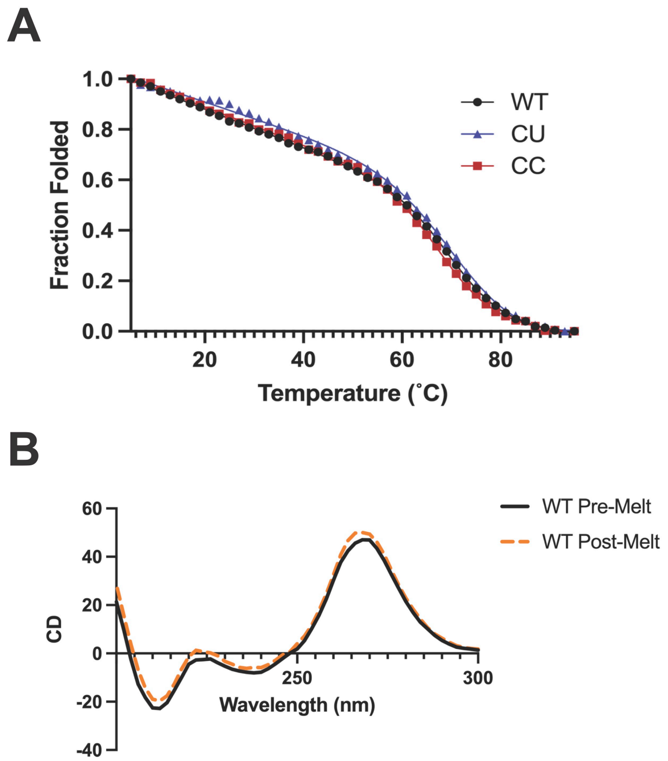


**Figure S8. CD unfolding of A-loop RNA constructs.** A) Full CD unfolding data from 5 to 95 °C. RNA constructs demonstrated cooperative two-state unfolding. B) CD wavelength scans from 300 to 200 nm of the WT A-loop construct before the melting experiment (black) and after the melting experiment (orange). The similarity of the curves implies reversible unfolding of the RNA during melting experiments.

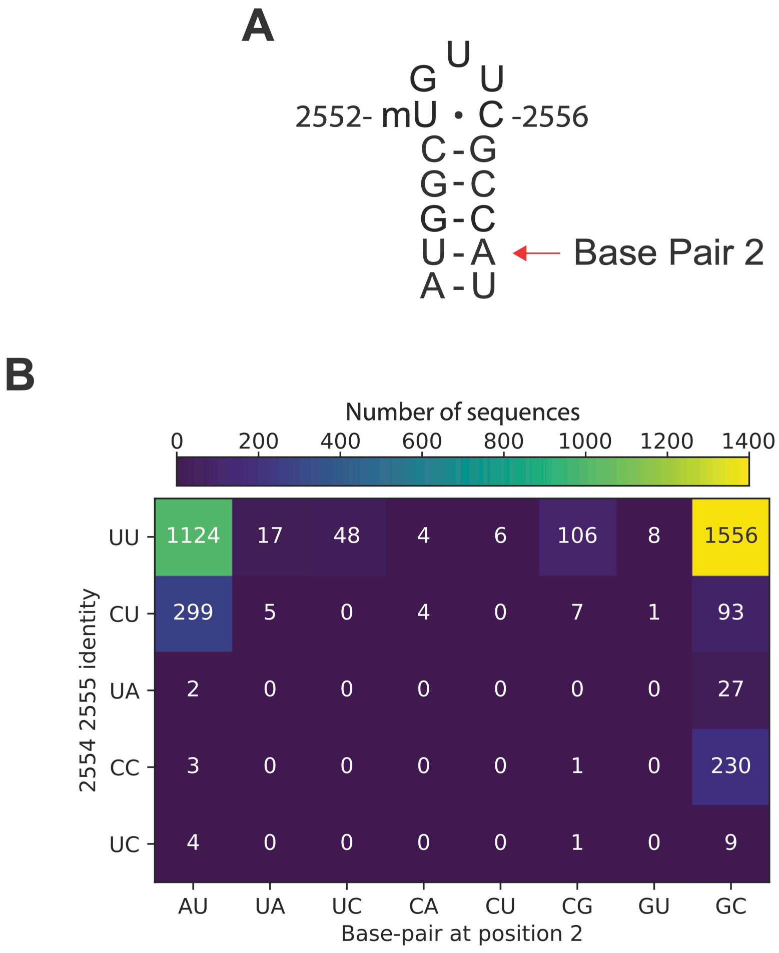


**Figure S9. Archaeal sequences at the base of their A-loop.** A) Base pair 2 (2548-2560) is highlighted on the secondary structure of the *E. coli* A-loop. B) Distribution of archaeal sequences in the SILVA database based on 2554/2555 and base pair 2 identity. The left axis classifies Archaea based on the identity of nucleotides at positions 2554 and 2555. The main populations are UU, CU, and CC. The bottom axis classifies Archaea based on the nucleotide identity at base pair 2 in the A-loop. Almost all Archaea with cytidines at positions 2554 and 2555 have a GC base pair at position 2. Around half of Archaea with uridines at positions 2554 and 2555 also have a GC base pair at position 2.

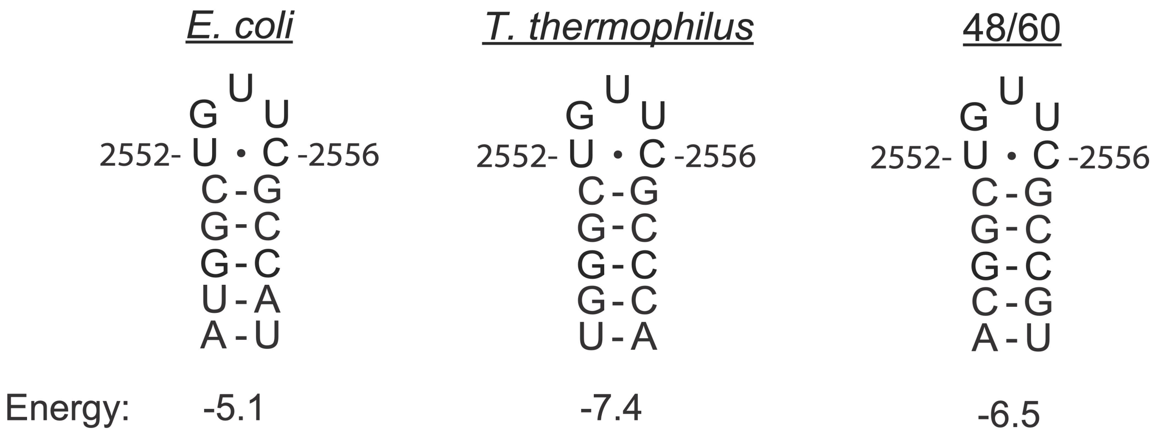


**Figure S10. Secondary structures and predicted minimum free energies of A-loop mutants.** *T. thermophilus* has variation at the base of the A-loop that stabilize its secondary structure. The 48/60 mutant was designed to stabilize the base of the A-loop while leaving pyrimidine bases at positions 2548 and 2561 to maintain contacts with ribosomal protein uL14. Relative minimum free energies were calculated in RNAstructrure.[2]

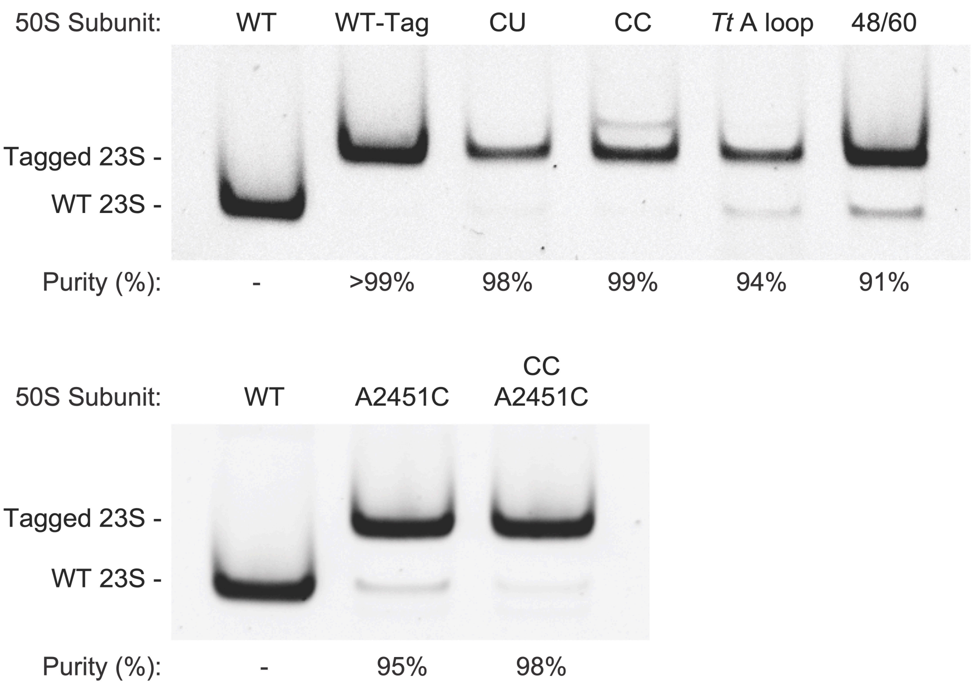


**Figure S11. Purity of 50S subunits used for *in vitro* assays.** After MS2 purification, 23S rRNA was isolated from 50S subunits and used for RT-PCR analysis. DNA containing an MS2 tag sequence is 32 bp larger than DNA with the native 23S sequence. RT-PCR products were run on a 10% polyacrylamide-TBE gel, and band intensities were used to quantify ribosome purity.

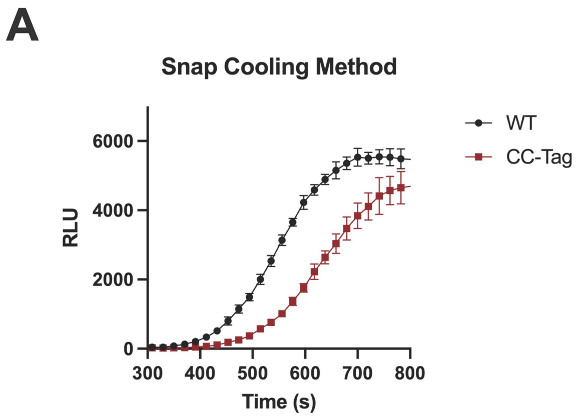


**Figure S12. HiBit translation assay for untagged WT and MS2-tagged CC 50S ribosomes.** Ribosomes were pre-incubated at 50 °C for 30 minutes and then cooled quickly on ice (left) or allowed to cool more slowly by incubating at room temperature for an additional 30 minutes (right).

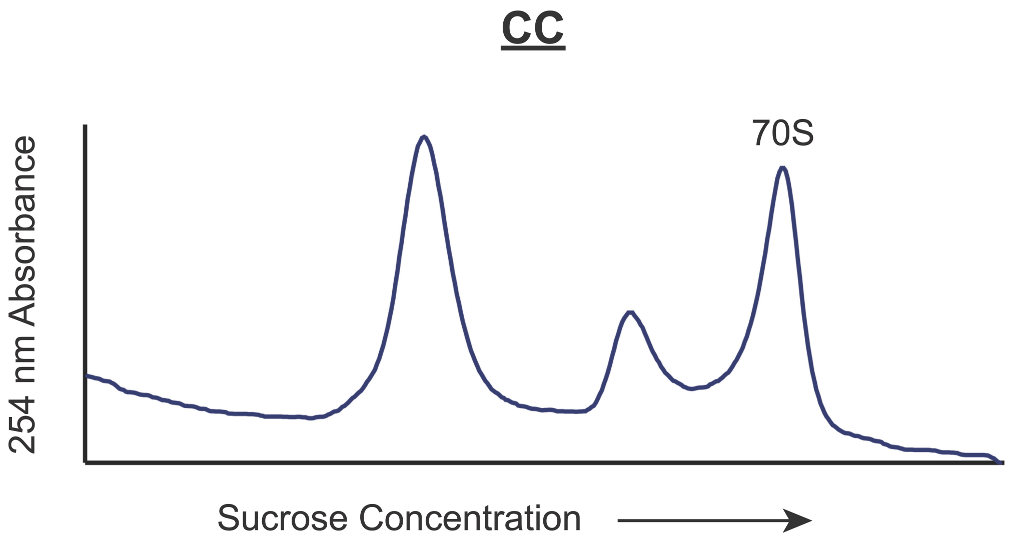


**Figure S13. Association gradient of CC mutant 50S and WT *E. coli* 30S subunits.** 30S subunits (2 equiv.) and 50S subunits (1equiv.) were incubated with 10 mM $MgCl_2$ and resolved on a 15-40% sucrose gradient. 70S fractions were collected for cryo-EM analysis.

**Table S1. Doubling times for *E. coli* expressing plasmid encoded ribosome mutants**

| pLK35 Plasmid Sequence | Doubling Time (h) |
|---|---|
| WT | 1.09 ± 0.01 |
| CU | 1.03 ± 0.01 |
| CC | 1.08 ± 0.02 |

**Table S2. Thermodynamic parameters derived from CD melting experiments for A-loop RNA constructs.**[+]

| Construct | $T_m$ (° C) | $\Delta H_{unfold}$ (kJ/mol) | $\Delta G_{25}$[++] (kJ/mol) |
|---|---|---|---|
| WT | 71.8 ± 0.6 | 197 ± 9 | 26.8 ± 0.9 |
| CU | 71.8 ± 0.1 | 189 ± 2 | 25.7 ± 0.3 |
| CC | 69.0 ± 0.2 | 183 ± 8 | 23.6 ± 1.2 |

+ Errors are reported as the S.D. of two experimental replicates.

++ $\Delta G_{25}$ is the Gibbs free energy of unfolding at 25 °C.

### Table S3. Additional Growth Temperatures for Archaeal Taxa

| Name | Strain | TaxID | Taxonomy | Growth Temperature | Source |
|---|---|---|---|---|---|
| Nitrosopelagicus brevis | CN25 | 1410606 | Archaea;Thaumarchaeota | 22 °C | [3] |
| Nitrosotalea devanaterra | - | 1078905 | Archaea;Thaumarchaeota | 25 °C | [4] |
| Nitrosotenuis aquarius | - | 1846278 | Archaea;Thaumarchaeota | 33 °C | [5] |
| Nitrosocaldus islandicus | - | 1846278 | Archaea;Thaumarchaeota; Nitrososphaeria | 65 °C | [6] |
| Nitrosocosmicus franklandus | C13 | 1798806 | Archaea;Thaumarchaeota; Nitrososphaeria; Nitrososphaerales; Nitrososphaeraceae | 37.5 °C | [7] |
| Caldiarchaeum subterraneum | - | 1798806 | Archaea;Thaumarchaeota | 70 °C | [8] |

**Table S4. DNA primers used in this study**

(m denotes 2'-O-Me; lowercases in the sequence indicate sites where mutations are introduced)

| Primer name | Sequence | Comment |
|---|---|---|
| U2554C_F | GGTATGGCTGcTCGCCATTTA | Forward primer for U2554C 23S rRNA mutagenesis |
| U2554C_R | CTTGGGACCTACTTCAGC | Reverse primer for U2554C 23S rRNA mutagenesis |
| U2554/5C_F | GGTATGGCTGccCGCCATTTAAAG | Forward primer for U2554C U2555C 23S rRNA mutagenesis |
| U2554/5C_R | CTTGGGACCTACTTCAGC | Reverse primer for U2554C U2555C 23S rRNA mutagenesis |
| Tt_Aloop_F | TCGCCcaTTAAAGTGGTACGCGAGC | Forward primer for A2547U U2548G A2560C U2561A 23S rRNA mutagenesis (*Tt* A loop) |
| Tt_Aloop_R | ACAGCCcaACCCTTGGGACCTACTTC | Reverse primer for A2547U U2548G A2560C U2561A 23S rRNA mutagenesis (*Tt* A loop) |
| 48/60_Aloop_F | TCGCCgTTTAAAGTGGTACGCGAG | Forward primer for U2548C A2560G 23S rRNA mutagenesis (48/60) |
| 48/60_Aloop_R | ACAGCCgTACCCTTGGGACCTACTT | Reverse primer for U2548C A2560G 23S rRNA mutagenesis (48/60) |
| A2451C_F | TCCGGGGATAcCAGGCTGATA | Forward primer for A2451C 23S rRNA mutagenesis |
| A2451C_R | GTACCTTTTATCCGTTGAGC | Reverse primer for A2451C 23S rRNA mutagenesis |
| MS2_quant_F | CTTGCCCCGAGATGAGTTCTCCC | RT-PCR primer for the ribosome purity assay |
| MS2_quant_R | GTACCGGTTAGCTCAACGCATCGCT | Primer for amplification of cDNA in the ribosome purity assay |
| HiBit Template | GCGAATTAATACGACTCACTATAGGGTTAACTTTAAC AAGGAGAAAAACATGGTGAGCGGCTGGCGCCTGTTT AAAAAAATTAGCTAACTAGCATAACCCCTCTCTAAAC GGAGGGGTTTAGTCA | DNA template for the HiBit peptide |
| HiBit_amp_F | GCGAATTAATACGACTCACTATAG | Forward amplification primer for the HiBit template |
| HiBit_amp_R | AAACCCCTCCGTTTAGAG | Reverse amplification primer for the HiBit template |
| $tRNA^{fMET}$ Template (C1G) | AATTCCTGCAGTAATACGACTCACTATAGGCGGGGT GGAGCAGCCTGGTAGCTCGTCGGGCTCATAACCCGA AGGTCGTCGGTTCAAATCCGGCCCCCGCAACC | DNA template for $tRNA^{fMet}$ C1G -A |
| fMet_amp_F | AATTCCTGCAGTAATACGACTCAC | Forward amplification primer for the $tRNA^{fMet}$ template |
| fMet-A_amp_R | mGmGTTGCGGGGGCC | Reverse amplification primer for the $tRNA^{fMet}$ template |

**Table S5. RNA oligos (IDT) used in this study.** (m denotes 2'-*O*-Me)

| **Primer name** | **Sequence** |
|---|---|
| WT A-loop | AUGGCmUGUUCGCCAU |
| CU A-loop | AUGGCmUGCUCGCCAU |
| CC A-loop | AUGGCmUGCCCGCCAU |

**Table S6. Cryo-EM Data Collection and Processing**

| | |
|---|---|
| Magnification | 105,000 |
| Voltage (kV) | 300 |
| Electron Exposure ($e^-$/Å$^2$) | 40 |
| Defocus Range (μm) | -0.5/-1.5 |
| Pixel Size (Å) | 0.8293 |
| Symmetry Imposed | C1 |
| Initial Particle Images | 1,614,362 |
| Final Particle Images | 114,493 |
| Map Resolution (Å) | 2.21 |
| FSC Threshold | 0.143 |

### Table S7. Model Refinement Statistics

| | |
|---|---|
| Model component | |
| Model resolution (Å) | 2.21 |
| FSC threshold | 0.143 |
| Map sharpening *B* factor (Å$^2$) | -38.8 |
| Model composition | |
| Non-hydrogen atoms | 148866 |
| $Mg^{2+}$ ions | 268 |
| $Zn^{2+}$ ions | 2 |
| Polyamines | 17 |
| Waters | 6756 |
| Ligands (paromomycin) | 1 |
| Mean *B* Factors (Å$^2$) | |
| RNA | 15.98 |
| Protein | 15.81 |
| Waters | 8.73 |
| Other | 10.56 |
| R.m.s. deviations from ideal values | |
| Bond (Å) | 0.005 |
| Angle (°) | 0.762 |
| Molprobity all-atom clash score | 8.93 |
| Ramachandran plot | |
| Favored (%) | 96.37 |
| Allowed (%) | 3.58 |
| Outliers (%) | 0.05 |
| RNA validation | |
| Angles outliers (%) | 0.004 |
| Sugar pucker outliers (%) | 0.2 |
| Average suiteness | 0.587 |

### References


1. Watson, Z. L. *et al.* All-atom metadynamics captures preferred geometries of non-L-a-amino acid monomers within the ribosomal PTC. *In preparation* (2022).
2. Bellaousov, S., Reuter, J. S., Seetin, M. G. & Mathews, D. H. RNAstructure: Web servers for RNA secondary structure prediction and analysis. *Nucleic Acids Res.* **41**, 471–474 (2013).
3. Santoro, A. E. *et al.* Genomic and proteomic characterization of 'Candidatus Nitrosopelagicus brevis': An ammonia-oxidizing archaeon from the open ocean. *Proc. Natl. Acad. Sci. U. S. A.* **112**, 1173–1178 (2015).
4. Lehtovirta-Morley, L. E., Stoecker, K., Vilcinskas, A., Prosser, J. I. & Nicol, G. W. Cultivation of an obligate acidophilic ammonia oxidizer from a nitrifying acid soil. *Proc. Natl. Acad. Sci. U. S. A.* **108**, 15892–15897 (2011).
5. Sauder, L. A., Engel, K., Lo, C., Chain, P. & Neufeld, J. D. "Candidatus Nitrosotenuis aquarius," an Ammonia-Oxidizing Archaeon from a Freshwater Aquarium Biofilte. *Appl. Enviromental Microbiol.* **84**, (2018).
6. Daebeler, A. *et al.* Cultivation and genomic analysis of 'Candidatus Nitrosocaldus islandicus,' an obligately thermophilic, ammonia-oxidizing thaumarchaeon from a hot spring biofilm in Graendalur valley, Iceland. *Front. Microbiol.* **9**, 1–16 (2018).
7. Lehtovirta-Morley, L. E. *et al.* Isolation of 'Candidatus Nitrosocosmicus franklandus', a novel ureolytic soil archaeal ammonia oxidiser with tolerance to high ammonia concentration. *FEMS Microbiol. Ecol.* **92**, 1–10 (2016).
8. Beam, J. P. *et al.* Ecophysiology of an uncultivated lineage of Aigarchaeota from an oxic, hot spring filamentous 'streamer' community. *ISME J.* **10**, 210–224 (2016).
